# The trajectory of gait development in mice

**DOI:** 10.1101/774885

**Authors:** Shyam Akula, Katherine B. McCullough, Claire Weichselbaum, Joseph D. Dougherty, Susan E. Maloney

## Abstract

Gait irregularities are prevalent in neurodevelopmental diseases and disorders (NDDs). However, there is a paucity of information on gait phenotypes in the NDD experimental models. This is in part due to the lack of understanding of the normal developmental trajectory of gait maturation in the mouse. Using the DigiGait system, we have developed a quantitative, standardized, and reproducible assay of developmental gait metrics in commonly used mouse strains that can be added to the battery of mouse model phenotyping. With this assay, we characterized the trajectory of gait in the developing C57BL/6J and FVB/AntJ mouse lines. In both lines, maturation of mouse gait between P21 and P30 was best reflected by aspects of the stance phase of the stride. Additionally, in C57BL/6J mice, gait maturation was observed through how the paw is loaded and unloaded during the stance phase of a stride. In FVB/AntJ mice, gait maturity during this developmental window was best observed through the decreased variability in paw area on the belt during stance. Our results also underscore the importance of considering body length when interpreting gait metrics. Further, our results show that background strain needs to be considered when comparing gait across models. Overall, our results show that aspects of mouse gait development parallel a timeline of normal human gait development, such as the percent of stride that is stance phase and swing phase. This study may be used as a standard reference for developmental gait phenotyping of a variety of murine models, such as models of neurodevelopmental disease.

## Introduction

Consistent gait is a marker of coordination and normal neurological function. Gait disturbances are the hallmark phenotype of diseases like cerebral palsy and Parkinsonism and can be observed in acute states of neurological dysfunction, such as alcohol intoxication (Nieuwboer et al., 2001; Vonghia et al., 2008; Wren et al., 2005). Subtle gait differences are also a feature of many neurodevelopmental disorders (NDDs), such as Autism Spectrum Disorders (ASD) and Williams Beuren Syndrome (WS) (Hocking et al., 2008; Kindregan et al., 2015). Gait disturbances in NDDs may be a consequence of underlying alterations in circuit function, or in circuit maturation. Mice are often used to study the function of normal circuits, their development, and their disruption in disease states. While the behavioral output of these circuits has been studied in adult animals, the maturation of gait has not been characterized in mice.

Understanding how the development of gait is affected in human NDDs requires an understanding of the healthy development of gait. The foundation for gait is thought to be reflexive and repetitive, as stepping patterns exhibited by infants are at least in part generated by the spinal cord (Forssberg, 1985). Maturation of gait into adult-like patterns requires coordinated activity of multiple systems including vestibular, musculoskeletal, and central nervous systems (CNS). Thus, it is likely the disruption of gait in NDDs reflects altered maturation of the CNS circuitry.

Gait is made up of strides that comprise a stance phase with the foot in contact with the ground and a swing phase with the foot off the ground. In children, walking begins around 12 - 14 months of age and a mature pattern of gait is established by around 3 years of age (Pediatric Musculoskeletal Matters International; Sutherland et al., 1980). By five to seven years old gait resembles that of an adult (Hillman et al., 2009; Lythgo et al., 2011; Sutherland et al., 1980). Adult gait is composed of a stride pattern generally made up of about 60% stance phase and 40% swing phase with a cycle involving a heel strike, stance and then toe propulsion. Children reach an adult pattern of gait through a decrease in double support time (both feet in stance simultaneously), and a decrease in stance duration with an increase in swing duration, suggesting a decrease in the stance/swing ratio with age. Markers of gait maturation also include a decrease in the number of strides per second (cadence) and increased stride length, yet these two metrics are driven by limb lengthening and greater limb stability (Sutherland et al., 1980) and may not reflect a maturation of the neural circuits underlying gait production.

Variability in gait, however, may better reflect refinement in the underlying neural circuitry. Motor control is known to arise from several factors including a reduction in noise within the sensorimotor system, and a refined organization and use of perceptual feedback within this system (Deutsch and Newell, 2005). An essential component of gait maturation is gaining adequate motor control such that variability is reduced (Adolph et al., 2015). Therefore, intraindividual variability serves as a proxy for motor control and, thus, maturation of central gait circuitry. In children, the variability in stride length, stance time, and double support time decreases with age, with substantial reductions during the first few months after onset of walking (Adolph et al., 2015). Decreases in variability continue until around 15 years of age, with particular changes observed in stride time variability from 3 to 14 years of age (Adolph et al., 2015). Thus, these reports suggest circuits underlying motor control are likely undergoing maturation through adolescence.

Gait has long been studied in the mature mouse, but the pattern of gait maturation in this experimental system is poorly defined. Thus, a comprehensive study of how gait develops including body size changes, spatiotemporal and posture parameters, and intraindividual variability is needed. Modern image and video analysis allow for computerized gait analysis systems that expand the quantifiable gait parameters to include temporal and postural metrics alongside the spatial metrics produced with footprint analysis on ink and paper. One such system is the DigiGait (Mouse Specifics, Boston, MA), a treadmill system with a transparent belt that allows creation of digital “footprints” to analyze posture and kinematics through capturing images of the mouse underside and paws. Leveraging this system, we can comprehensively define spatiotemporal and postural aspects of gait, as well as the intraindividual variability within these metrics. Further, we can identify which metrics are influenced by changing body size, and thus would be less interesting from the point of view of studying circuits. Studying how gait develops will enable us to better understand how behavioral motor circuits are refined and matured, and, thus, guide future studies into abnormalities in circuit function and maturation in NDD.

To this end, we characterized normative gait in two inbred mouse strains, C57BL/6J and FVB/AntJ, across development using the DigiGait gait analysis system. Our assay begins at postnatal day (P)21, the youngest age at which the mice could reliably complete the treadmill assay, and an age that corresponds to approximately 2-3 years of age in humans in terms of brain development (Gegenhuber and Tollkuhn, 2019; Semple et al., 2013). The assay continued through the juvenile stage, covering a window of time during which substantial maturation occurs in human gait (Pediatric Musculoskeletal Matters International; Sutherland et al., 1980). We present below how spatiotemporal and postural gait metrics and their intraindividual variability change with and without the influence of body size. These data provide a detailed examination of gait maturation in the mouse. Further, they provide an index which can inform interpretations of future studies of altered gait development in mouse models of disease.

## Materials and Methods

### Animals

All experimental protocols were approved by and performed in accordance with the relevant guidelines and regulations of the Institutional Animal Care and Use Committee of Washington University in St. Louis. C57BL/6J (C57; https://www.jax.org/strain/000664) and FVB/AntJ (FVB; https://www.jax.org/strain/004828) wildtype inbred strains were used in this study. This FVB substrain is homozygous for the wild-type *Pde6b* allele and thus does not go blind. Twenty-six (7 M, 19 F) C57 and 32 (15 M, 17 F) FVB mice were used. To limit variability observed in gait studies conducted in a cross-sectional design (Hillman et al., 2009), we performed a longitudinal study. This resulted in a reduced sample size for C57 mice to 9 (5 M, 4 F) and FVB mice to 19 (7 M, 12 F) due to a reduced number of quantifiable videos from these mice at all time points. All mice used in this study were maintained and bred in the vivarium at Washington University in St. Louis. For all experiments, adequate measures were taken to minimize any pain or discomfort. The colony room lighting was 12:12h light/dark cycle; room temperature (∼20-22°C) and relative humidity (50%) controlled automatically. Standard lab diet and water was freely available. Pregnant dams were individually housed in translucent plastic cages measuring 28.5×17.5×12cm with corncob bedding. Upon weaning at postnatal day (P)21, mice for behavioral testing were group housed according to sex.

### Gait Analysis

#### Apparatus

Gait data was collected using the DigiGait Imaging System (Mouse Specifics, Inc), an advanced gait analysis system with Ventral Plane Imaging Technology that generates digital paw prints from the animal as it runs on a motorized treadmill (Hampton et al., 2004). This system has been described in detail previously (Hampton et al., 2004). Briefly, the floor of the apparatus was made of tempered glass around which looped a transparent belt (PCV and HDPE blend) measuring 6 cm wide. The animal was sequestered to the 40.5 cm run alley by a transparent polycarbonate enclosure and illuminated to 1381 lux from above and below with LED panel lighting. A digital video camera (Basler Ace/Scout Camera with 12.5/16mm lens) was housed beneath the run alley to capture the ventral plane image of the animal, which was sent to the DigiGait software on a connected computer. Due to smaller body size of juvenile mice, 0.6 cm thick bumpers made from expanded PVC were placed on the sides of the apparatus using magnets to narrow the width of the run alley and thus increase the potential for straight, usable runs.

#### Procedure

Both male and female experimenters collected C57 gait data and the female experimenter collected FVB gait data. Pilot testing revealed FVB mice can adequately run on the DigiGait by P17. However, C57 mice could not run adequately until P21. Therefore we chose to examine gait across the post-weaning, juvenile period (Figure 1A). All testing occurred during the light phase of the circadian cycle. For all habituation and testing sessions, the mice were placed in a holding room adjacent to the testing environment for a 30 min acclimation period prior to testing, and weighed. Prior to testing the paws of the FVB mice were dyed with red food coloring (McCormick) diluted into water to increase contrast with belly fur. The testing room overhead lights were off during testing, however the holding room lights remained on to decrease disruptions of circadian rhythms. On P20, each mouse was habituated to the apparatus. This consisted of placing the animal on the stationary belt and starting the belt moving at 5 cm/s and slowly increasing the speed until 20 cm/s is reached allowing for at least 30 sec of run time. Testing occurred P21, P24, P27 and P30. For these test days, each mouse was placed individually on the apparatus. The belt was started at 10 cm/s until the animal started walking forward. Once the animal reached the front of the alley, the speed was increased to 20 cm/s. Once a 3-5 sec usable run was acquired, the belt was stopped and the animal removed to the homecage. Criteria for a usable run included a consistent forward movement with no sliding, jumping, or side drift. The belt was cleaned with 70% EtOH between litters or as needed between mice and daily upon completion of testing.

**Figure 1.**
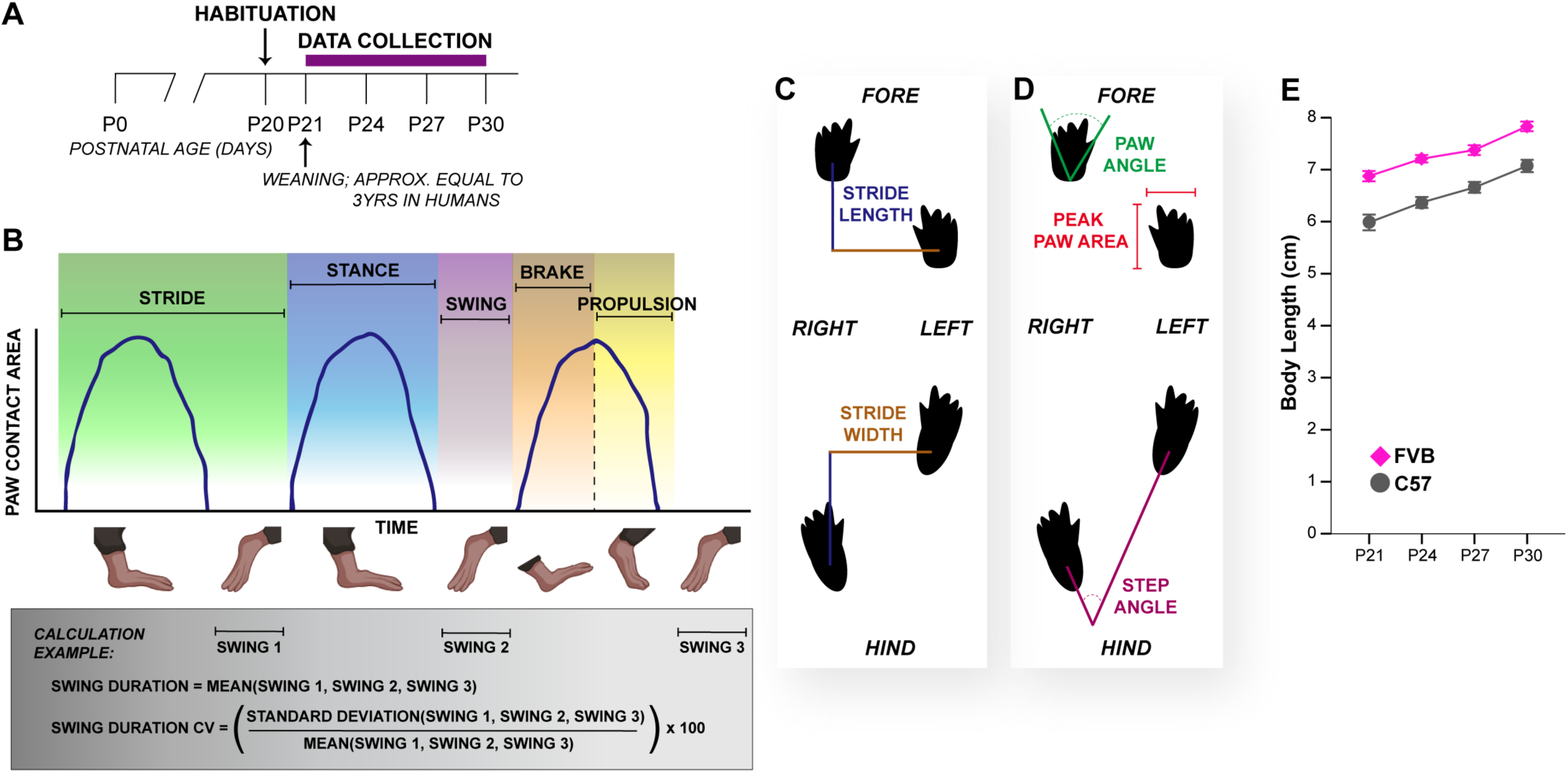
Gait analysis procedure and measurement schematics. (**A**) Schematic of developmental gait data collection procedure. Purple bar represents duration of data collection. (**B**) Schematic of paw contact area plots (blue lines) derived by DigiGait software to quantify spatiotemporal gait metrics (represented by different background colors). Below the graph is a cartoon representation of mouse feet during three strides. The gray box provides an example of variable calculations based on these plots. (**C**) Cartoon of digital mouse footprints with representations of measurements of the spatial metrics stride length (blue) and stride width (brown) measurements. (**D**) Cartoon of digital mouse footprints with representations of measurements of the postural metrics paw angle (green), step angle (eggplant), and peak paw area (red). (**E**) Body length measurements for C57 and FVB mice made along the long axis of the mouse from nose to base of tail (data are means ± SEM).

#### Video Processing

To obtain optimal contrast between the mouse paws and coat color for recognition of paws for digital footprint generation and processing of videos through the DigiGait software pipeline, we post-processed the videos using ImageJ (Wayne Rasband, NIH, USA). To determine optimal levels of brightness and contrast we used two videos from each time point and processed each multiple times using differing levels of brightness and contrast. The settings that resulted in the smoothest paw area contact plots were used (brightness adjustment of 19 and contrast adjustment of 135). These adjustments were then applied to all videos. The adjusted videos were exported as uncompressed .avi files to ensure compatibility with the DigiGait software.

Each video was then processed through the DigiGait Analysis software, as described previously (Hampton et al., 2004). Briefly, this entailed applying filters to exclude the snout and adjust contrast for optimal digital footprint generation. Paw contact area plots were generated and validated against the video of the mouse. Any tracking errors were manually corrected. The digital footprints and paw contact area plots were then used to extract the gait parameters used for analysis (Figure 1B, described in detail below).

#### Selection of gait metrics for analysis

The DigiGait software analysis system outputs a comprehensive list of gait metrics. A concern with such a large list of variables is inflation of familywise error rate. To identify possible redundant variables, we generated scatterplot matrices. Pairs of variables that were visibly perfectly aligned were considered redundant and one variable within the pair was excluded from further analysis. The variables we excluded were stride duration, % brake of stride, % brake of stance, % propulsion of stride, % propulsion of stance, swing/stance ratio (Figures S1, S2), hindlimb shared stance time (Figures S3, S4), and ataxia coefficient (Figures S5, S6)

In addition to excluding variables based on redundancy, we also determined if variables required exclusion based on poor reliability. The post-video acquisition processing within the DigiGait software requires adjustment of filters to remove the snout and decrease noise from the digital paw prints, as well as manual corrections to errors within the paw contact area plots, introducing the possibility of inconsistencies across experimenters. We examined the inter-rater reliability of this processing between the measurements produced by two independent experimenters by calculating intraclass correlation coefficients (ICC) with their 95% confidence intervals using IBM SPSS Statistic software (v.25) based on absolute-agreement and two-way mixed-effects model. The data used for this were derived from an independent cohort of 10 FVB mice tested on P24, P27 and P30. For fore and hind limbs, respectively, 20/25 and 27/32 metrics showed excellent reliability (ICC ≥ .75) and another 2/25 and 2/32 showed good reliability (ICC = .60 - .74). The remaining metrics showed poor reliability (ICC ≤ .39; 3/25 and 3/32). ICCs are reported in Table 1. We excluded from further analysis any metrics with poor reliability: midline distance, axis distance, paw drag, and maximal rate of change of paw area contact during the propulsion phase. We also excluded tau propulsion despite good reliability because we felt this measure has not yet been defined or validated adequately in the literature and thus its usefulness is not clear.

**Table 1.** Gait metrics organized by subtype with definitions and intraclass correlation coefficients (ICC) with their 95% confidence intervals used to determine inter-rater reliability of gait video processing between the measurements produced by two independent experimenters.

| Component Subtype and Metric |  | Definition |  | ICC with 95% CI |  |
| --- | --- | --- | --- | --- | --- |
| Spatial Subcomponent |  |  |  | Fore | Hind |
|  | Stride Frequency | Number of completed stides per second (cadence). |  | .975 [.958, .985] | .989 [.981, .993] |
|  | Stride Length | Distance covered during one full 'rotation' of a paw through both stance and swing phases. |  | .986 [.976, .992] | .991 [.985, .995] |
|  | Stance Width | Distance between fore or hind limbs during full stance. |  | .972 [.940, .987] | .990 [.978, .995] |
|  | Overlap Distance | Average overlapping distance of ipsilateral paws across successive strides. |  | n/a | .994 [.991, .997] |
|  | Paw Placement Positioning | The extent of overlap of ipsilateral paws at full stance (reflecting balance). |  | n/a | .989 [.981, .994] |
|  | Gait Symmetry | The ratio of left to right step frequency. |  | n/a | .610 [.336, .771] |
| Temporal Subcomponent |  |  |  |  |  |
|  | Swing Duration | Time (ms) the paw is not in contact with the belt. |  | .972 [.952, .983] | .982 [.970, .990] |
|  | Stance Duration | Time (ms) the paw is in contact with the belt. |  | .985 [.974, .991] | .995 [.991, .997] |
|  | Brake Duration | Time of the braking portion of the stance phase where the paw is initiating contact with the belt though the heel (initial paw contact to full paw contact; immediately follows swing phase). |  | .946 [.908, .968] | .932 [.883, .960] |
|  | Propulsion Duration | Time of the propelling portion of the stance phase where the paw is lifting off of the belt though the toes (full paw contact to final paw contact; immediately precedes swing phase). |  | .954 [.921, .973] | .980 [.966, .988] |
|  | Stance Factor | The ratio of left to right stance durations (measure of gait symmetry). |  | .949 [.890, .976] | .932 [.810, .972] |
|  | Maximal rate of paw contact change | Maximal rate of paw area contact change during the braking portion of stance (how quickly the paw is loaded on to the belt). |  | .838 [.724, .905] | .775 [.617, .868] |
|  | % Stance | Percent of stride that comprises the stance phase. |  | .971 [.950, .983] | .981 [.967, .989] |
|  | % Swing | Percent of stride that comprises the swing phase. |  | .971 [.950, .983] | .981 [.967, .989] |
|  | % Hindlimb Shared Stance Time | Percent of stance phase during which both hind limbs are in contact with the belt. |  | n/a | .982 [.967, .990] |
| Postural Subcomponent |  |  |  |  |  |
|  | AbsolutePawAngle | The angle of the paw with the long axis of the direction of locomotion of the animal (degree of external rotation). |  | .909 [.845, .947] | .948 [.911, .969] |
|  | StepAngle | The angle between the right and left hind paws due to stride length and stance width. |  | .898 [.982, .953] | .995 [.987, .998] |
|  | PeakPawArea | Area of the paw at full stance. |  | .751 [.577, .853] | .783 [.631, .873] |
| Intraindividual Variability Parameters |  |  |  |  |  |
|  | Coefficient of Variance (CV) | A normalized measure of variability calculated as [(standard deviation/mean)x100]. |  | -- | -- |
|  | Stride Length CV |  |  | .875 [.787, .927] | .912 [.849, .949] |
|  | Stance Width CV |  |  | .960 [.914, .981] | .935 [.895, .970] |
|  | Swing Duration CV |  |  | .905 [.837, .944] | .909 [.844, .946] |
|  | Paw Angle CV |  |  | .853 [.749, .915] | .830 [.709, .900] |
|  | Step Angle CV |  |  | .671 [.283, .848] | .927 [.843, .966] |
|  | Peak Paw Area CV |  |  | .660 [.419, .800] | .980 [.975, .988] |

### Body Length Quantification

Animal body length was measured by importing videos from the Digigait software into Ethovision (Noldus Information Technology, The Netherlands). The animal’s body was automatically detected using the contrast of the darker body against a lighter background. A custom script provided by Noldus was used to calculate its length based on the coordinates of the nose, center point and tail base. To validate this method of body length measurement, the body lengths of a subset of FVB and C57 mice were also measured manually from the same videos, and intraclass correlation coefficients (ICC) with their 95% confidence intervals were calculated between the manual and automated measurements using IBM SPSS Statistics software (v.25) based on absolute-agreement and two-way mixed-effects model. The ICCs indicate excellent reliability between the manual and automated measurements (FVB, ICC = .972, 95% CI [.904, .992]; C57, ICC = .977, 95% CI [.955, .988]), providing confidence in our automated process for body length measurement.

### Statistical Analysis

All statistical analyses were performed using IBM SPSS Statistics software (v.25). Prior to analyses, all data were screened for missing values, fit of distributions with assumptions underlying univariate analyses. This included the Shapiro-Wilk test on *z*-score-transformed data and qqplot investigations for normality, Levene’s test for homogeneity of variance, and boxplot and *z*-scores (±3.29) investigation for identification of influential outliers. However, no outliers were removed. Means and standard errors were computed for each measure. Repeated measures analysis of variance (rmANOVA) was used to analyze gait data across juvenile ages. Where appropriate, the Greenhouse-Geisser or Huynh-Feldt adjustment was used to protect against violations of sphericity. The linear trend analysis contrast was used to examine linear change across ages and comparisons of P21 to P30 were conducted, both *a priori* determined. Post hoc comparisons across all ages were conducted and Bonferroni corrected following significant effects of age. Statistical results were confirmed with the non-parametric Friedman Test for any outcome measure with violations of normality. Violations of homogeneity of variance were rendered innocuous by equal sample sizes in ANOVA testing. To examine the influence of body length on gait metrics across age, linear mixed modeling (LMM) was used with age as a repeated fixed factor grouped by subject ID and body length as a covariate. The critical alpha value for all analyses was *p* < .05, unless otherwise stated, and *p* values were adjusted appropriately, as stated above. Test statistics and other details for each analysis are provided in Tables 2 and 3.

**Table 2.**
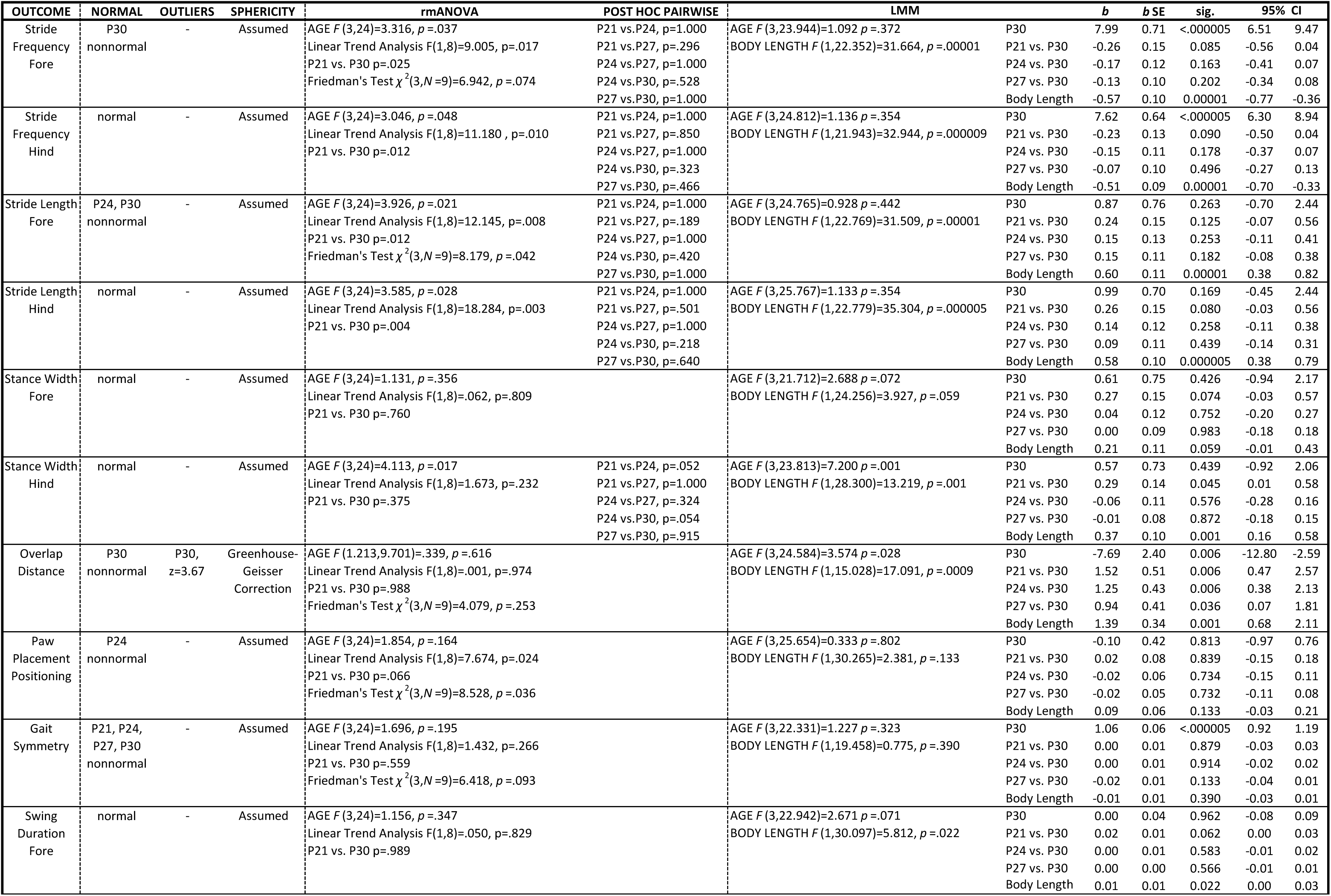
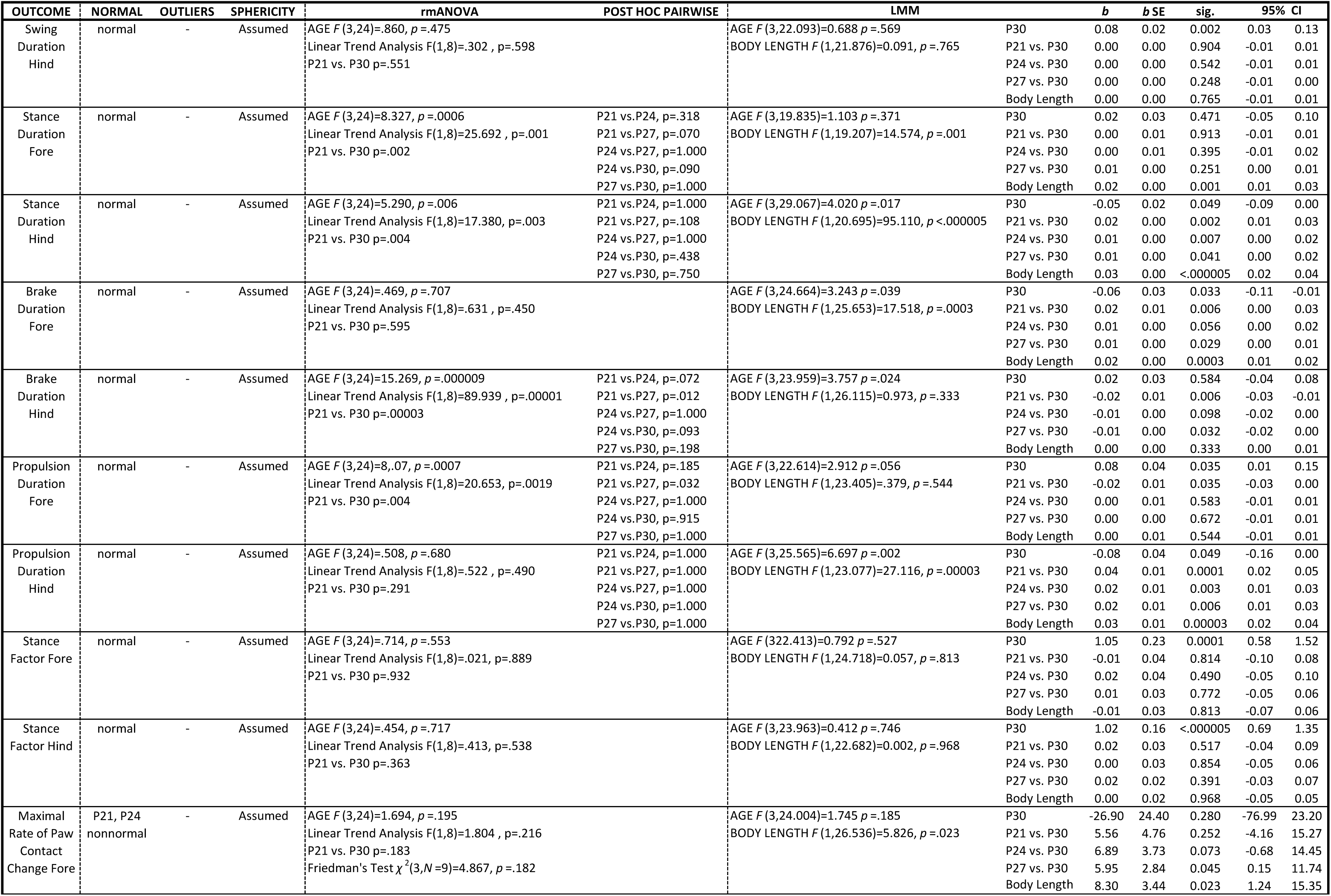
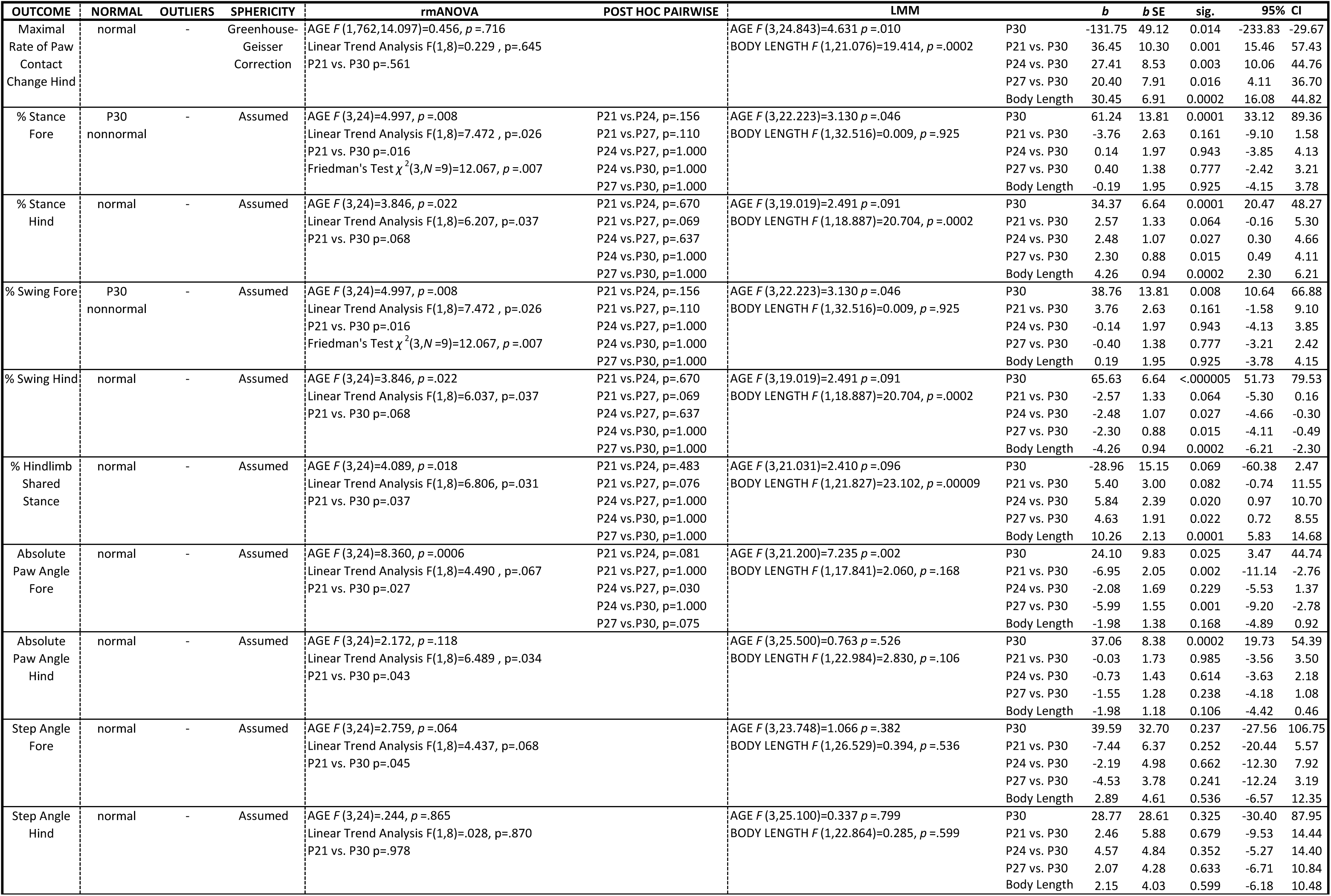
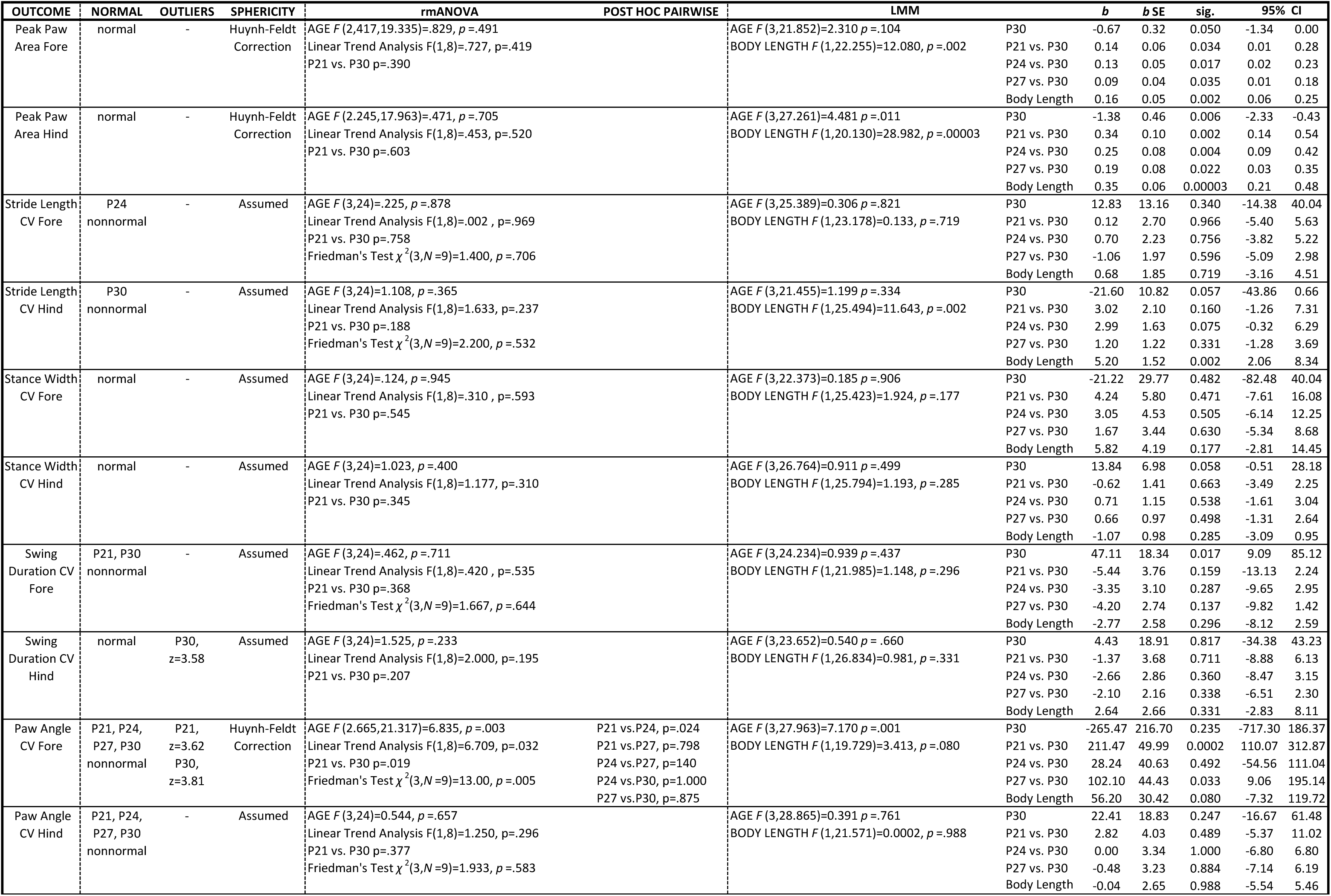
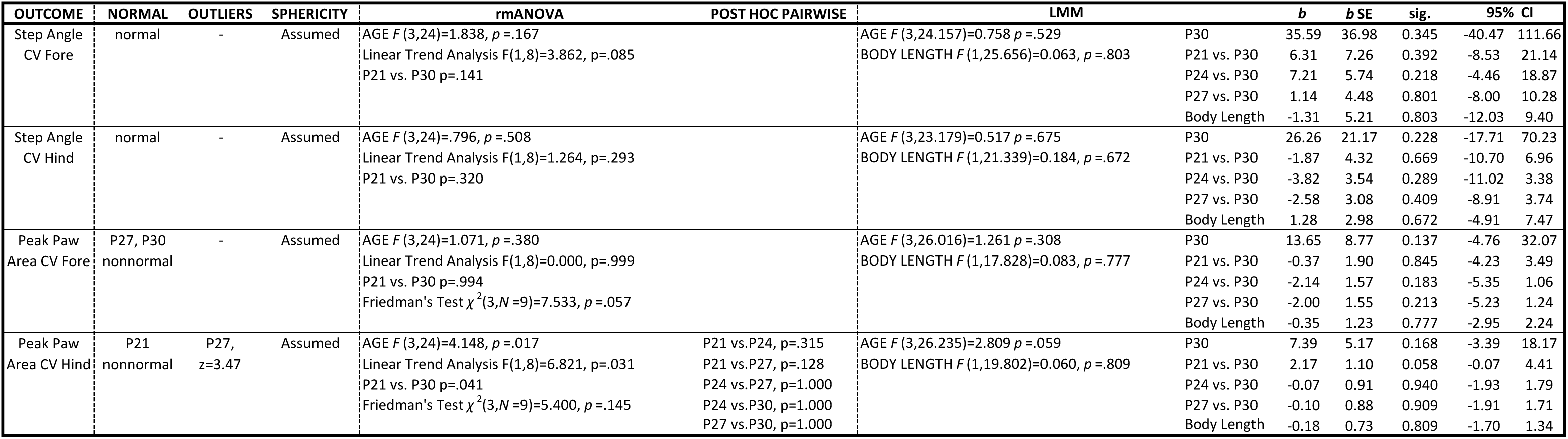
Statistical output for C57 gait data.

**Table 3.**
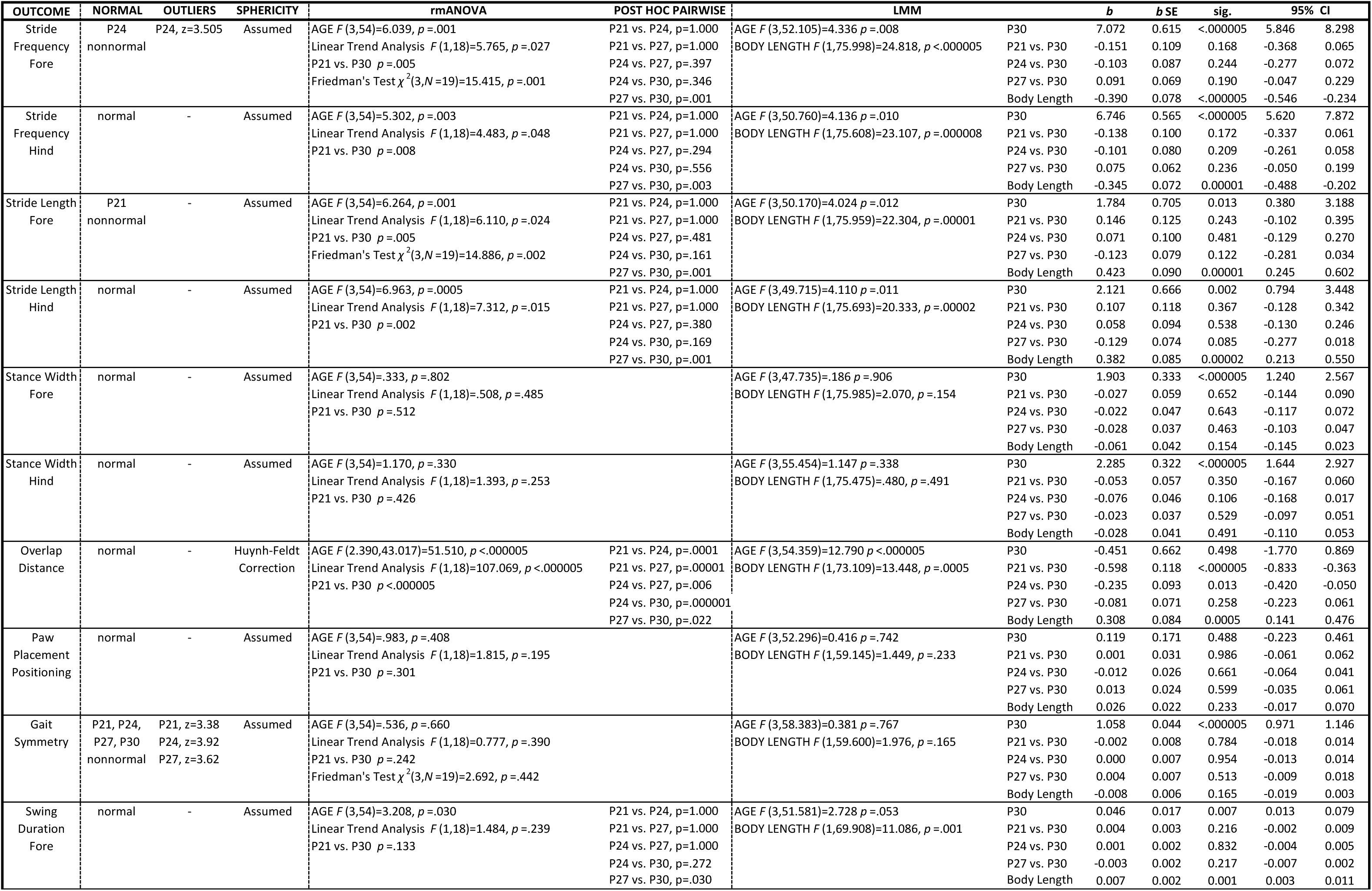
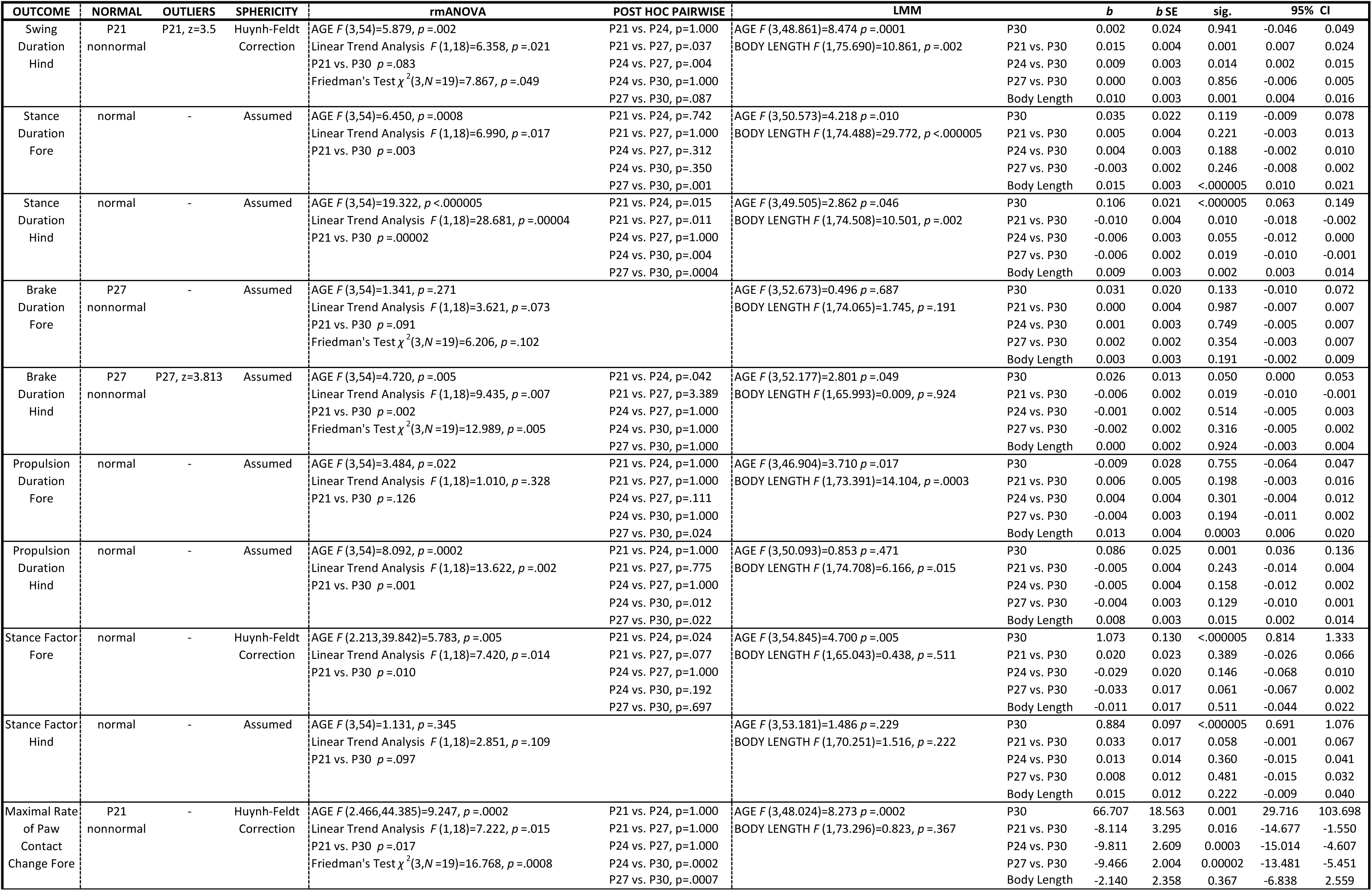
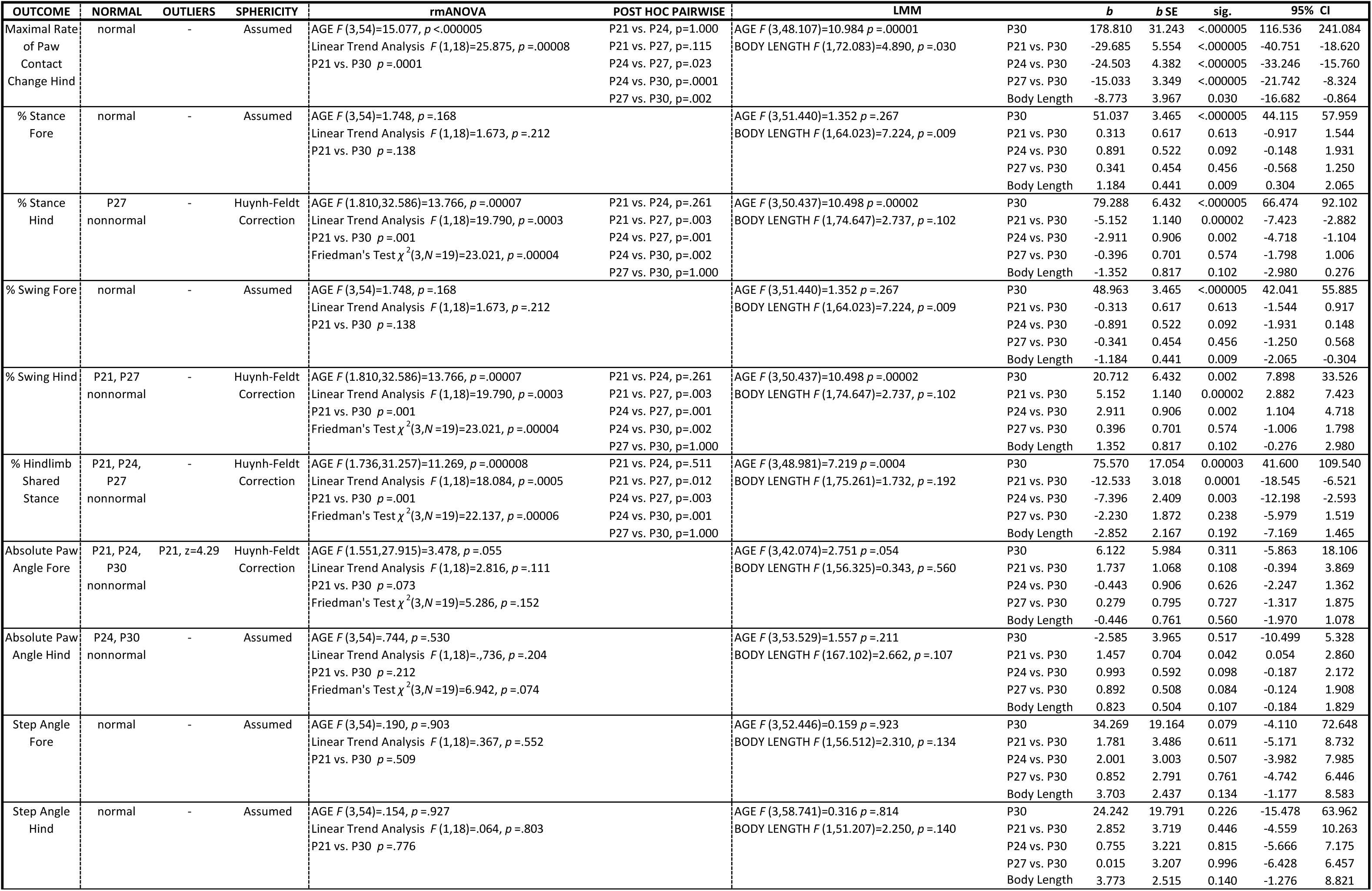
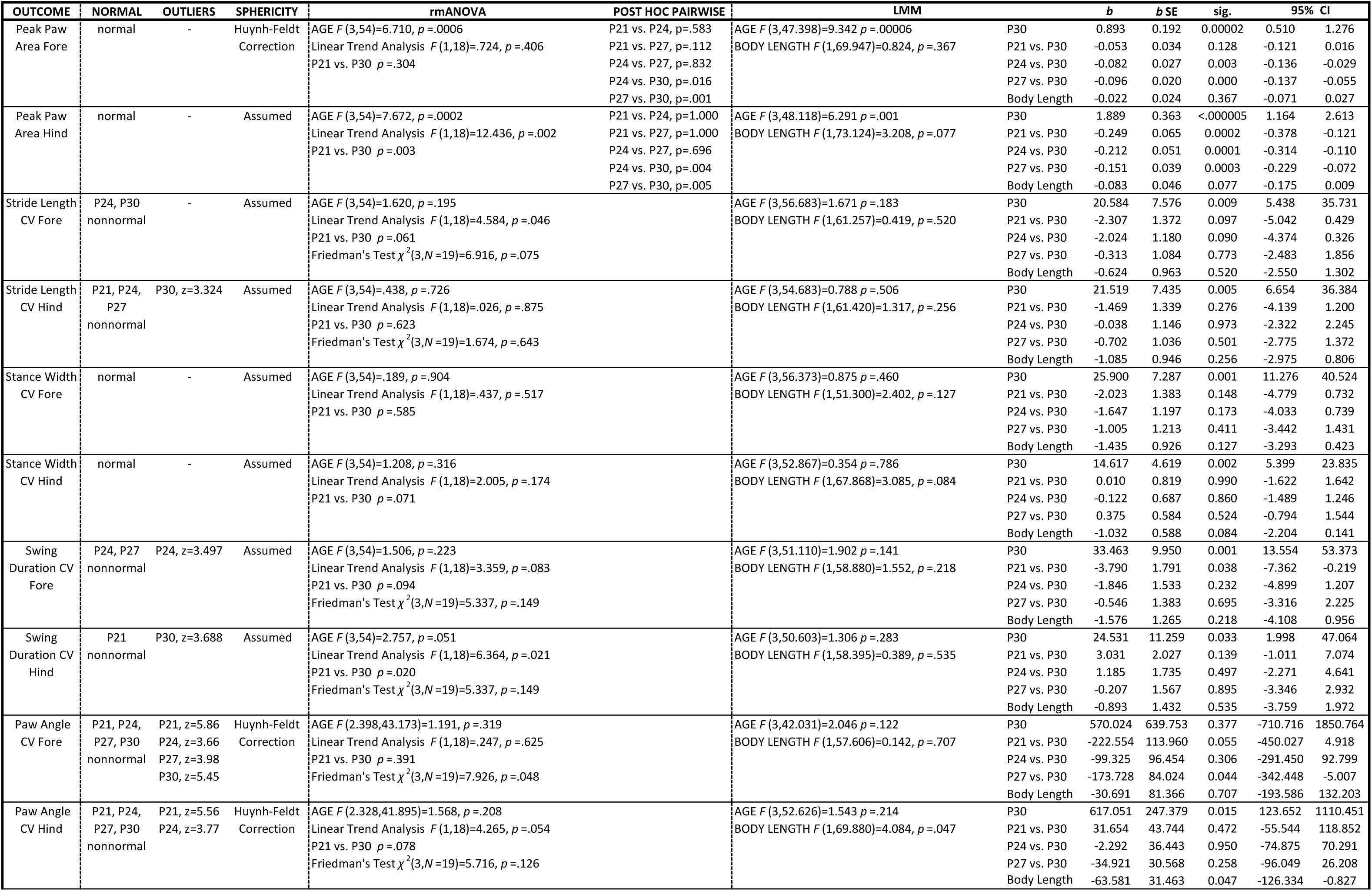
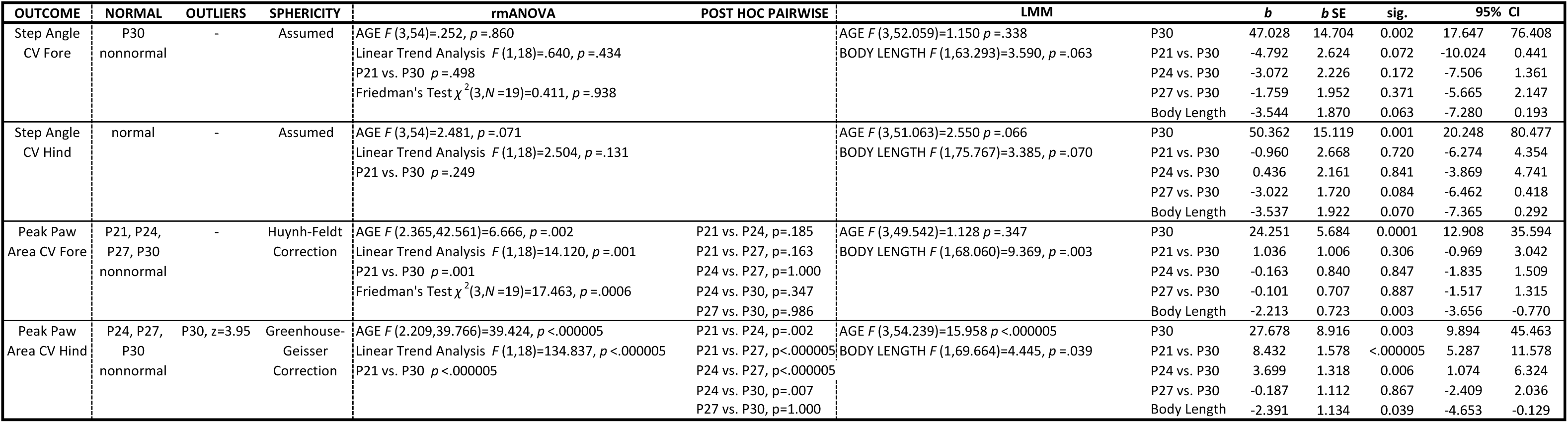
Statistical output for FVB gait data.

### Data Availability Statement

The datasets generated and analyzed during the current study are available from the corresponding author upon reasonable request.

## Results

Gait is an automatic motor behavior, and while it can be voluntarily modified, is normally produced unconsciously in a stereotyped manner. Gait is usually analyzed by quantifying components of the step cycle, or stride, broken into when a paw has contact with the ground, known as the stance phase, and when it is moving through the air, known as the swing phase (Figure 1B). The stance phase is further broken down into the paw braking phase (heel strike to full stance) and propulsion phase (full stance to toe push off). Using the DigiGait, distinct gait metrics can be quantified (Table 1), including spatiotemporal measures such as stride length, stance width, and propulsion duration (Figure 1B,C), and postural measures such as step angle, paw angle, and peak paw area during full stance (Table 1, Figure 1D). The DigiGait software extracts the temporal measures from the paw contact area plots derived from the digital footprints (Figure 1B), while the spatial and postural measures are derived straight from the digital footprints (Figure 1C,D). Each of these measures is calculated as an average across all strides of a trial (See Figure 1B for example). All our trials comprised at least 12 strides based on previous work suggesting 9 strides or more are required for DigiGait data processing (Hampton et al., 2004) (C57, *M*=19.0, range: 12.0-24.5; FVB, *M*=18.9, range: 13.0-25.5). The intraindividual variability within many of the measures was also calculated as the coefficient of variance (CV) by dividing the standard deviation of the strides in a trial by the mean of the strides in a trial (See Figure 1B for example). Thus, we were able to examine meaningful gait metrics across development that reflect the maturation of spatiotemporal, postural, and variability metrics.

In our study design, we addressed multiple issues that might confound our interpretations of our gait measurements. A major consideration for our study is the accompanying change in body length across the age range examined (Figure 1E,F) and how that increase may influence gait. To account for this, we conducted a second set of analyses using LMM to adjust for body length measurements across age by using it as a covariate. This allowed us to interpret the effect of age on each gait parameter while controlling for the influence of body length. Both statistical models, rmANOVA and LMM with the covariate, are presented below to identify metrics that are dependent and those that are independent of body length. To reduce variability that can result from cross-sectional designs in gait studies, we limited our analysis to longitudinally collected data. Finally, because speed is the greatest influencer of gait, the speed of the treadmill during data collection was kept constant across all ages at 20 cm/s to allow for appropriate comparisons of forced gait across age.

### Gait metrics were heavily impacted by body length changes in C57 mice

We found that many gait metrics significantly changed from P21 to P30 (Table 4) in C57 mice, allowing us to identify likely gait maturation patterns across this developmental window. These metrics represented all four subcomponents of gait, with the majority reflecting changes to temporal dynamics; all are discussed in detail below. However, body length significantly influenced 43% of the gait variables we examined in C57 mice, substantially affecting our interpretation of gait changes during this developmental window. Results from the rmANOVA suggested 17 of 44 metrics significantly changed from P21 to P30 (Table 2). However, after adjusting for the influence of body length in the LMM, this was reduced to 12 metrics, only 7 of which overlapped with the original 17. Thus, our data presented below highlight the importance of considering body length when interpreting gait performance in the mouse.

**Table 4.**
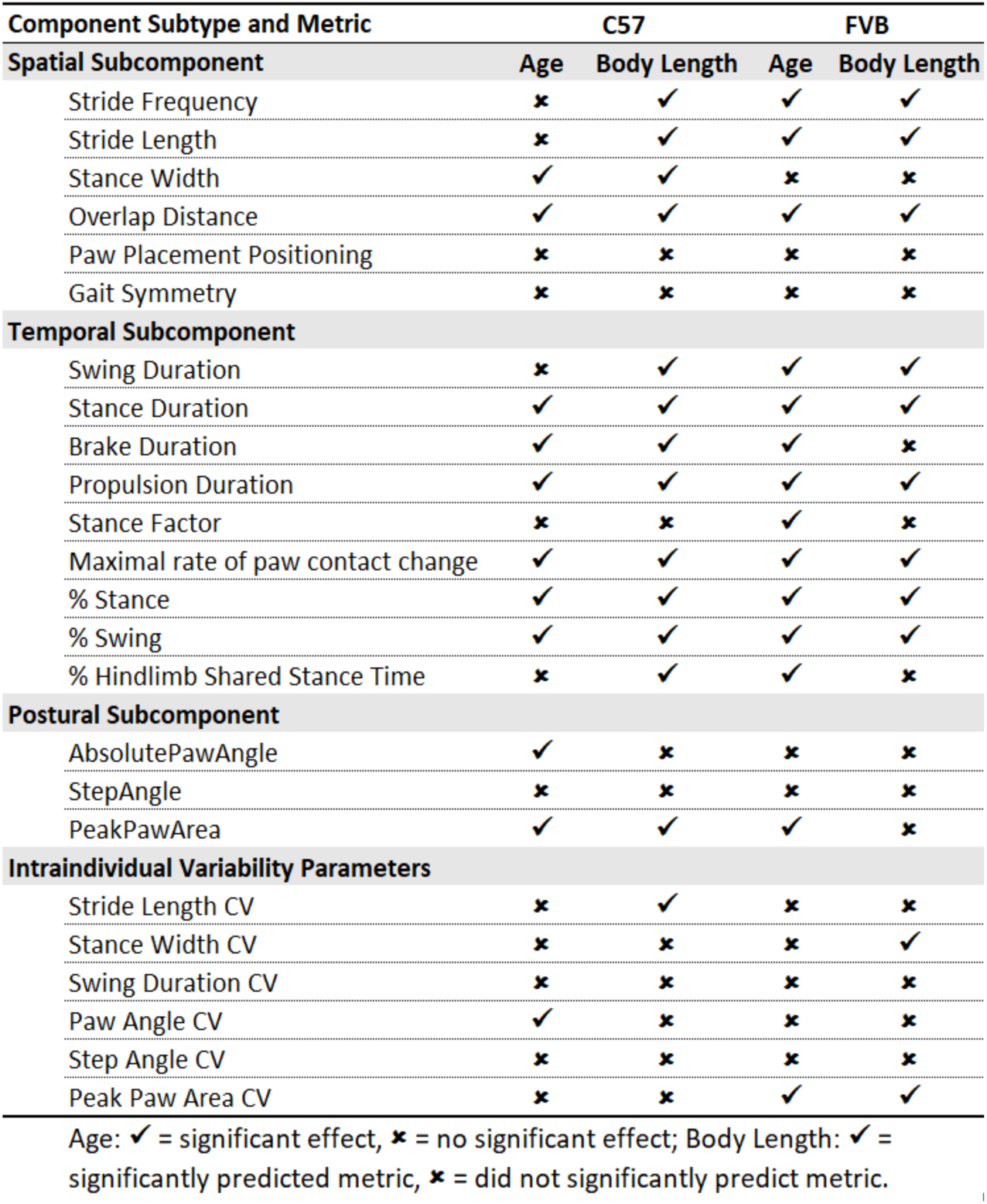
Chart of Age and Body Length influence on gait metrics measured across P21 to P30 in C57 and FVB mice.

| Component Subtype and Metric | C57 |  | FVB |  |
| --- | --- | --- | --- | --- |
| Spatial Subcomponent | Age | Body Length | Age | Body Length |
| Stride Frequency | ✗ | ✓ | ✓ | ✓ |
| Stride Length | ✗ | ✓ | ✓ | ✓ |
| Stance Width | ✓ | ✓ | ✗ | ✗ |
| Overlap Distance | ✓ | ✓ | ✓ | ✓ |
| Paw Placement Positioning | ✗ | ✗ | ✗ | ✗ |
| Gait Symmetry | ✗ | ✗ | ✗ | ✗ |
| Temporal Subcomponent |  |  |  |  |
| Swing Duration | ✗ | ✓ | ✓ | ✓ |
| Stance Duration | ✓ | ✓ | ✓ | ✓ |
| Brake Duration | ✓ | ✓ | ✓ | ✗ |
| Propulsion Duration | ✓ | ✓ | ✓ | ✓ |
| Stance Factor | ✗ | ✗ | ✓ | ✗ |
| Maximal rate of paw contact change | ✓ | ✓ | ✓ | ✓ |
| % Stance | ✓ | ✓ | ✓ | ✓ |
| % Swing | ✓ | ✓ | ✓ | ✓ |
| % Hindlimb Shared Stance Time | ✗ | ✓ | ✓ | ✗ |
| Postural Subcomponent |  |  |  |  |
| AbsolutePawAngle | ✓ | ✗ | ✗ | ✗ |
| StepAngle | ✗ | ✗ | ✗ | ✗ |
| PeakPawArea | ✓ | ✓ | ✓ | ✗ |
| Intraindividual Variability Parameters |  |  |  |  |
| Stride Length CV | ✗ | ✓ | ✗ | ✗ |
| Stance Width CV | ✗ | ✗ | ✗ | ✓ |
| Swing Duration CV | ✗ | ✗ | ✗ | ✗ |
| Paw Angle CV | ✓ | ✗ | ✗ | ✗ |
| Step Angle CV | ✗ | ✗ | ✗ | ✗ |
| Peak Paw Area CV | ✗ | ✗ | ✓ | ✓ |

Age: ✓ = significant effect, ✗ = no significant effect; Body Length: ✓ = significantly predicted metric, ✗ = did not significantly predict metric.

Excellent examples of the importance of body length in gait analysis were found with the metrics representing two main components of gait, stride frequency and stride length. These variables appeared to increase from P21 to P30 for both the fore- and hindlimbs (Figures 2A,B; Table 2). However, this change was entirely due to increased body length (Figure 1E). Body length was a significant predictor of both stride frequency and stride length, and when we regressed out the influence of body length, we no longer observed a significant change over time for either variable (Figures 2C,D). Body length also was a significant predictor of hindlimb, but not forelimb, stride length variability (CV), an intra-step variability metric (Figure 2E,F). However, this variability remained static across development regardless of the influence of body length. These data suggest that stride frequency and stride length (relative to body size) are established at an early age and do not reflect further maturation of gait during this developmental period.

**Figure 2.**
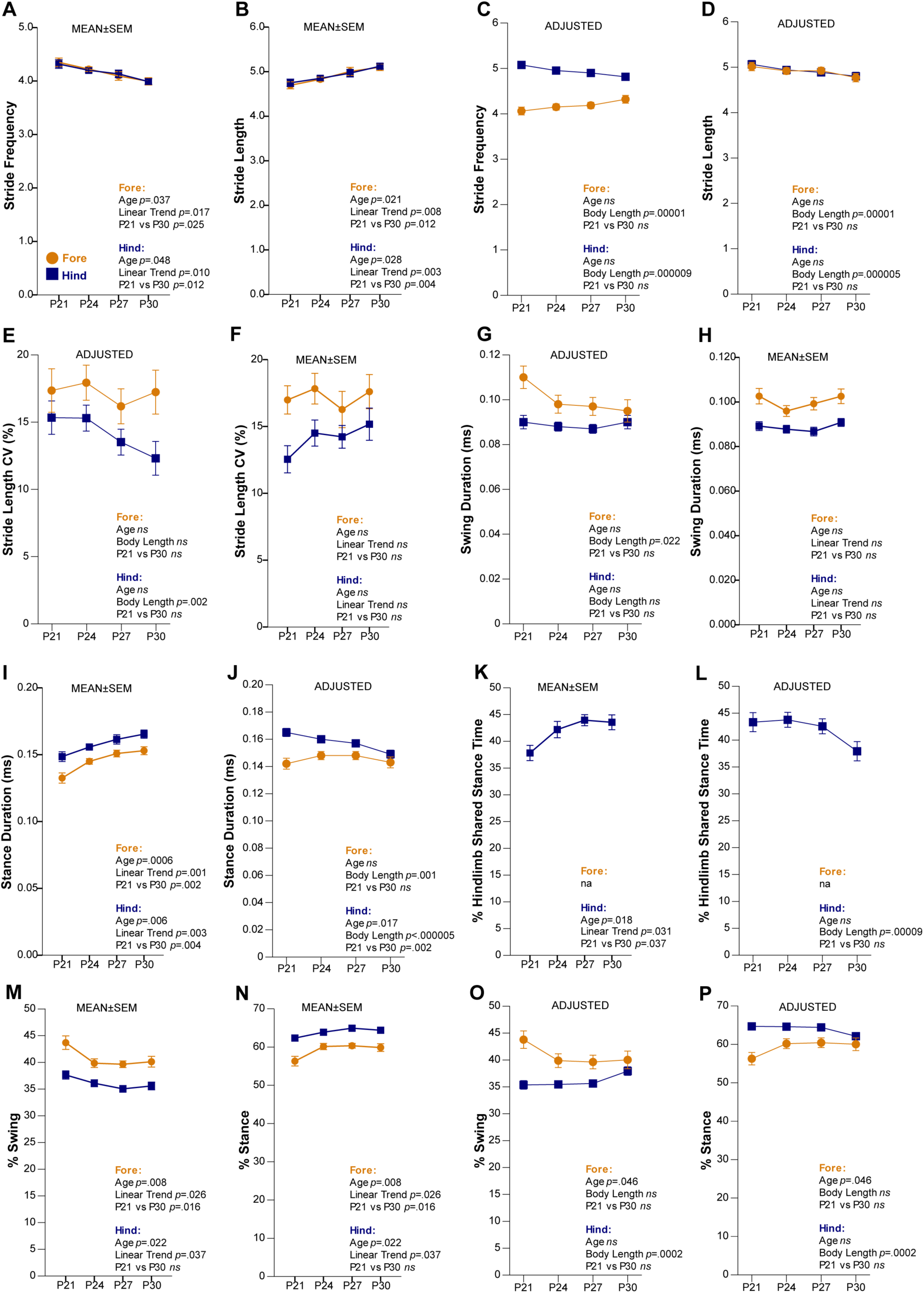
Main components of stride were significantly predicted by body length in C57 mice from P21 to P30. (**A-B**) Stride frequency and stride length means ± standard error of the mean (SEM). (**C-D**) Stride frequency and stride length covariate adjusted means (ADJUSTED) for body length (value 6.52078). (**E-F**) Stride length coefficient of variance (CV) means ± SEM and ADJUSTED. (**G-H**) Swing duration ADJUSTED and means ± SEM. (**I-J**) Stance duration means ± SEM and ADJUSTED. (**K-L**) %Hindlimb Shared Stance Time means ± SEM and ADJUSTED. (**M-P**) Means ± SEM and ADJUSTED for %swing and %stance. For means ± SEM graphs, statistics derived from rmANOVAs include main effect of Age, Linear Trend Analysis, and *a priori* determined pairwise comparison of P21 to P30. For ADJUSTED graphs, statistics derived from LMM include main effect of Age, main effect of the covariate Body Length and *a priori* determined pairwise comparison of P21 to P30. A comprehensive list of statistical output can be found in Table 2.

Next we examined the two main temporal components of a stride: the swing and stance phases. The swing duration of the forelimbs, but not hindlimbs, was predicted by body length and this variable was unchanged across development for both limbs (Figures 2G,H). For stance duration of fore- and hindlimbs the body length was a significant predictor. Across age, the stance duration for both the fore- and hindlimbs appeared to increase when body length was not covaried (Figure 2I). However, once this metric was adjusted for body length, forelimb stance duration no longer changed across the developmental window and hindlimb stance duration significantly decreased (Figure 2J). A related spatiotemporal measurement, %hindlimb shared stance time, also referred to as %double support time, also showed a significant increase across age when body length was not considered (Figure 2K). However, the results from the LMM revealed that the percent of the stride that consists of hindlimbs in shared stance remains constant across this developmental window (Figure 2L).

We also examined the percent of stride made up of the swing and stance phases, because changes to these metrics characterize gait maturation in humans (Sutherland et al., 1980). Without considering body length, both limbs appear to significantly decrease in %swing and increase in %stance across age (Figure 2M,N). For hindlimbs only, body length significantly predicted %swing and %stance components of stride (Figure 2O,P). Thus for forelimbs, the LMM confirmed the significant change over time for each %swing and %stance, while for the hindlimbs these metrics were unchanged across time. At P30, %swing for both limbs is ∼40% and %stance for both limbs is ∼60%, which is exactly the ratio observed in the mature human stride (Sutherland et al., 1980).

Spatial components of gait are also significantly predicted by body length and show developmental change. These include stance width (Figure 1C) and paw overlap distance (Table 1). The width of stance during gait appears not to change with age based on the rmANOVA results (Figure 3A). However, body length significantly predicted stance width and we saw that the stance width between the hindlimbs actually decreases when this metric is adjusted for body length in the LMM (Figure 3B). Another spatial metric, paw overlap distance, was heavily influenced by body length. This metric appeared unchanged in the rmANOVA (Figure 3C) yet showed a substantial decreasing slope from P21 - P30 when body length was considered in the LMM (Figure 3D). The decrease in distance between overlapping footprints of subsequent ipsilateral fore and hind paws may suggest greater flexibility of hip rotations or limb strength to achieve longer strides.

**Figure 3.**
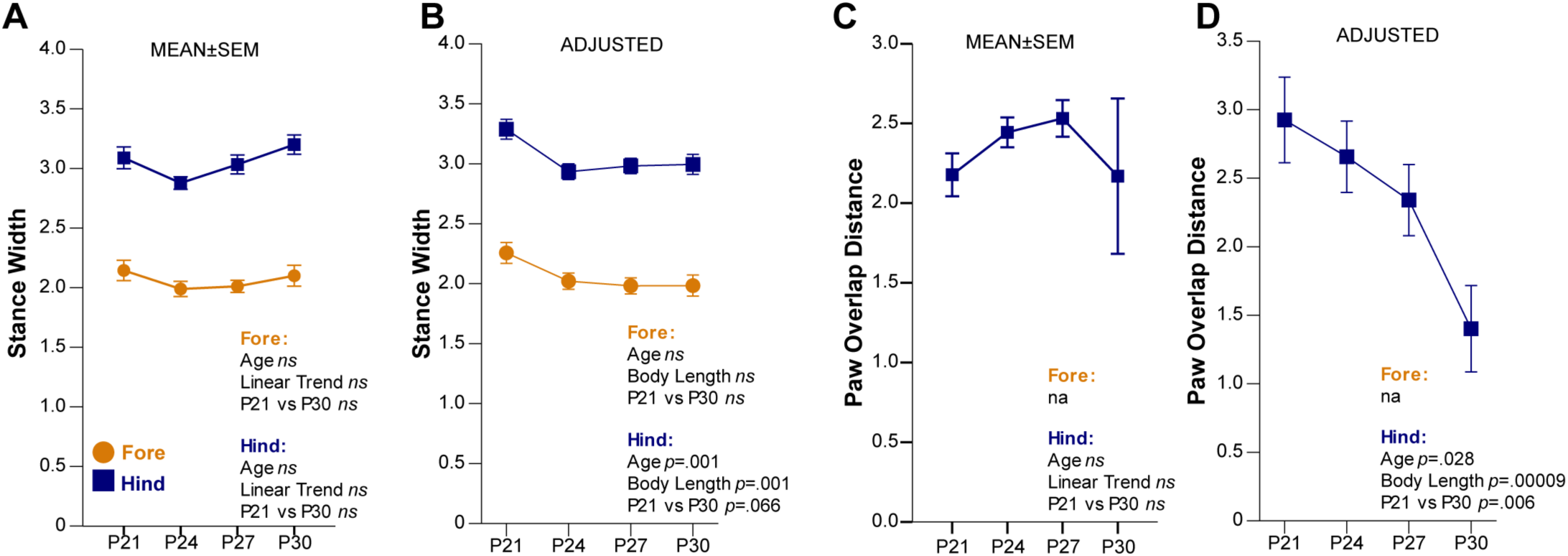
Distinct spatial gait metrics, when adjusted for the influence of body length, decreased from P21 to P30. (**A-B**) Stance width means ± standard error of the mean (SEM) and covariate adjusted means (ADJUSTED) for body length (value 6.52078). (**C-D**) Paw overlap distance means ± SEM and ADJUSTED. For means ± SEM graphs, statistics derived from rmANOVAs include main effect of Age, Linear Trend Analysis, and *a priori* determined pairwise comparison of P21 to P30. For ADJUSTED graphs, statistics derived from LMM include main effect of Age, main effect of the covariate Body Length and *a priori* determined pairwise comparison of P21 to P30. A comprehensive list of statistical output can be found in Table 2.

### Gait maturation in C57 mice was reflected by components of the stance phase

The metrics that appeared to best reflect gait maturation in C57 mice were spatiotemporal metrics that represent how the paw is loaded and unloaded from the stance phase of a stride. Specifically, the braking time and the propulsion time showed opposite developmental patterns for both fore- and hindlimbs in the LMM. The time the forelimbs were involved in braking decreased while the time the forelimbs were involved in propulsion remained consistent across time (Figures 4A,B). In contrast, the time the hindlimbs were involved in braking increased while the time the hindlimbs were involved in prolusion decreased. By P30, both sets of limbs were equally contributing to braking and propulsion. This balancing of the use of both fore- and hindlimbs for propulsion and braking may be an important aspect of gait maturation in the C57 mouse. Body length was a significant predictor of both braking and propulsion metrics, so the inclusion of body length in the model as a covariate changed the outcome. Specifically, the rmANOVA results suggested, in contrast to the LMM results, that the forelimbs braking time was unchanged while propulsion time increased with age, while in the hindlimbs the decrease in propulsion duration was not observed (Figures 4C,D). Body length also was a significant predictor of the maximal rate of paw contact change, or how quickly the paw is loaded onto the belt. Thus adding body length as a covariate allowed us to observe changes that were not revealed in the rmANOVA (Figure 4E,F). The rate at which the hindpaws were loaded onto the belt decreased across time, while that for the forelimbs remained unchanged. The slowing speed at which the hindpaw is loaded onto the belt is likely a major driver of the increased duration of the braking phase.

**Figure 4.**
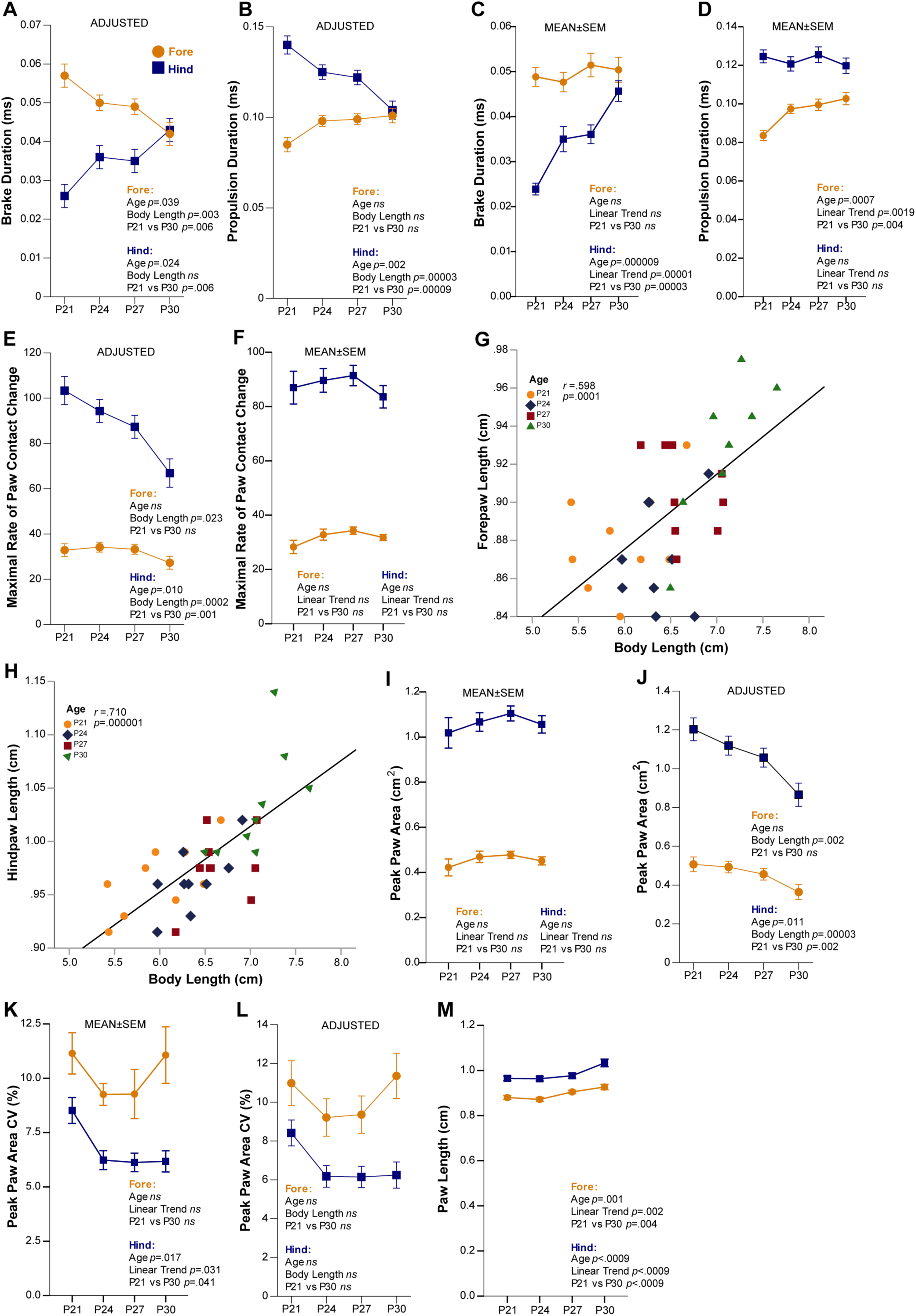
Gait metrics that represent the dynamics of the stance phase matured from P21 to P30 in C57 mice. (**A-D**) Brake and propulsion duration covariate adjusted means (ADJUSTED) for body length (value 6.52078) and means ± standard error of the mean (SEM). (**E-F**) Maximal rate of paw contact change ADJUSTED and means ± SEM. (**G-H**) Scatterplots of the relationship between body length and fore- and hindpaw lengths from P21 to P30. (**I-L**) Peak paw area and peak paw area coefficient of variance (CV) means ± SEM and ADJUSTED. (**M**) Paw length means ± SEM measured at full stance. For means ± SEM graphs, statistics derived from rmANOVAs include main effect of Age, Linear Trend Analysis, and *a priori* determined pairwise comparison of P21 to P30. For ADJUSTED graphs, statistics derived from LMM include main effect of Age, main effect of the covariate Body Length and *a priori* determined pairwise comparison of P21 to P30. A comprehensive list of statistical output can be found in Table 2.

Another metric related to the stance phase of the stride is peak paw area, which is measured at full stance. Peak paw area at full stance was significantly predicted by body length, which is expected because paw length is highly correlated with body length (Figures 4G,H). While no difference was observed in the rmANOVA for this metric without considering body length (Figure 4I), when body length was a covariate in the LMM a decrease in peak paw area for the hindlimbs was observed (Figure 4J). The variability of the peak paw area (peak paw area CV) remained unchanged across time (Figure 4K,L), and was independent of body length. The decreases across age to hindlimb maximal rate of paw contact change and hindlimb peak paw area may reflect a maturing of paw placement on the belt.

These two metrics, maximal rate of paw change and peak paw area, are likely also related to the paw size. Thus it is possible that the decreases in both metrics are reflections of a relatively larger size of the paws at younger ages compared to the body length. That is, the ratio of paw size to body length may decrease with age. To determine if this is the case, we measured paw lengths. We found both fore and hind paw lengths increased with age (Figure 4M) and that paw size is very strongly significantly correlated with body length (Figure 4I,J). Therefore, we believe the change in maximal rate of paw change and peak paw area is related to how the mice are loading their paws onto the belt, possibly reflecting a change towards heel-to-toe stepping from flatfooted stepping (Kraan et al., 2017).

The remainder of the metrics analyzed were independent of both body length and age, including the remaining postural subcomponents, aspects of gait variability, and measurements of gait symmetry. The postural metric absolute paw angle and its variability, and paw angle CV, did show significant changes across the developmental window measured for the forelimbs only, but the patterns are hard to interpret (Figures 5A-D). These metrics will need to be explored further to better elucidate their patterns across this age period. The postural metric, step angle and its companion step angle CV, were not influenced by body length nor did either change across ages (Figure 5E-H). In addition, the final measurements of variability to report, swing duration CV and stance width CV, were also not significantly influenced by body length or age (Figure 5I-L). These data suggest that maturation of these gait metrics occurs prior to P21. Lastly, the spatiotemporal metrics that reflect gait symmetry and balance were fully independent of body length and age (gait symmetry, stance factor, and paw placement positioning, Figure 6A-F). Maturation of these symmetry and balance variables likely occurs at very early ages, and it is unsurprising that we see no change in them at ages where the mice can already successfully run on a treadmill.

**Figure 5.**
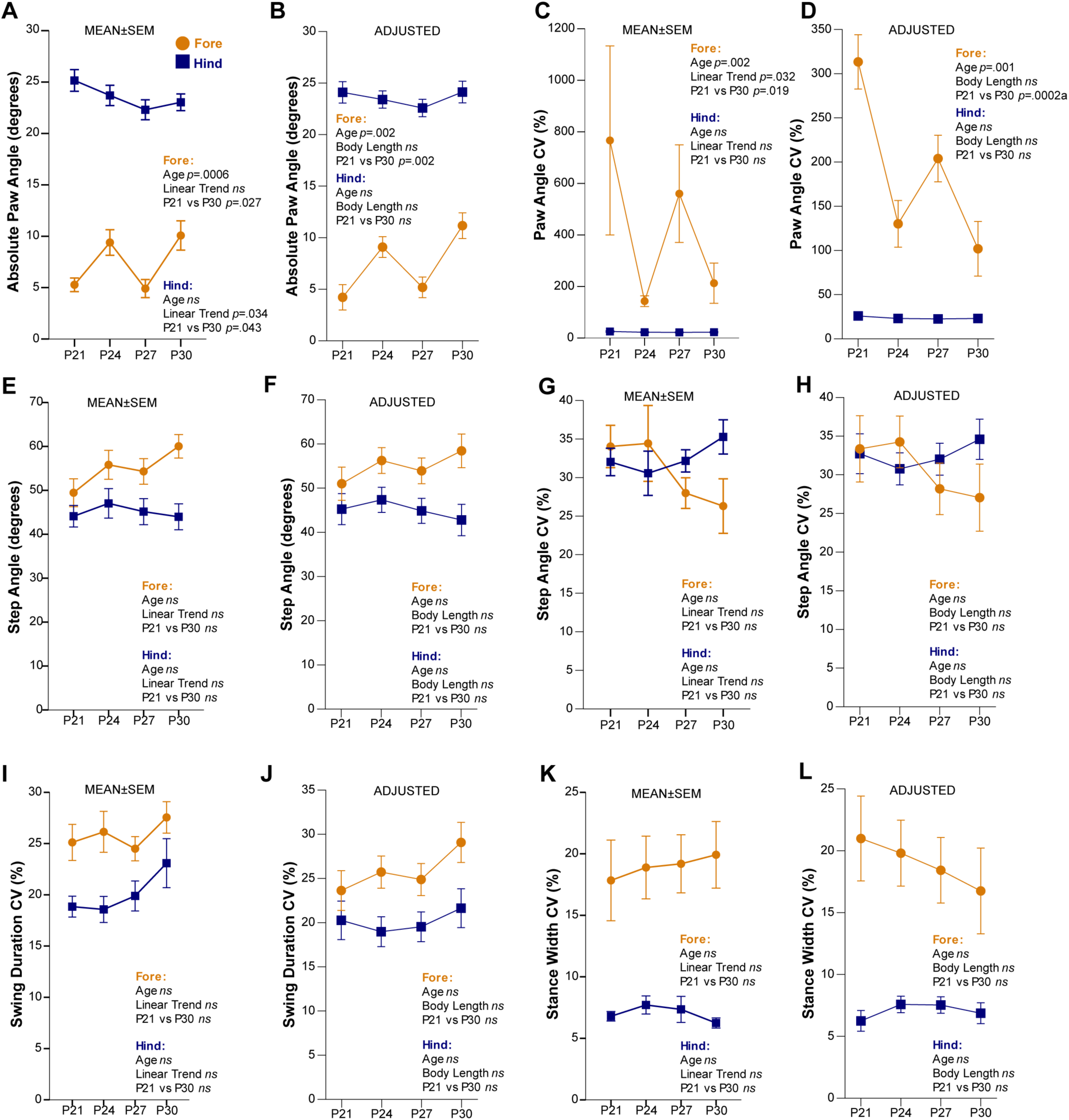
Postural and variability metrics measured from P21 to P30 in C57 mice were independent of both body length and age. (**A-D**) Absolute paw angle and paw angle coefficient of variance (CV) means ± standard error of the mean (SEM) and covariate adjusted means (ADJUSTED) for body length (value 6.52078). (**E-H**) Step angle and step angle CV means ± SEM and ADJUSTED. **(I-L**) Swing duration CV and stance width CV means ± SEM and ADJUSTED. For means ± SEM graphs, statistics derived from rmANOVAs include main effect of Age, Linear Trend Analysis, and *a priori* determined pairwise comparison of P21 to P30. For ADJUSTED graphs, statistics derived from LMM include main effect of Age, main effect of the covariate Body Length and *a priori* determined pairwise comparison of P21 to P30. A comprehensive list of statistical output can be found in Table 2.

**Figure 6.**
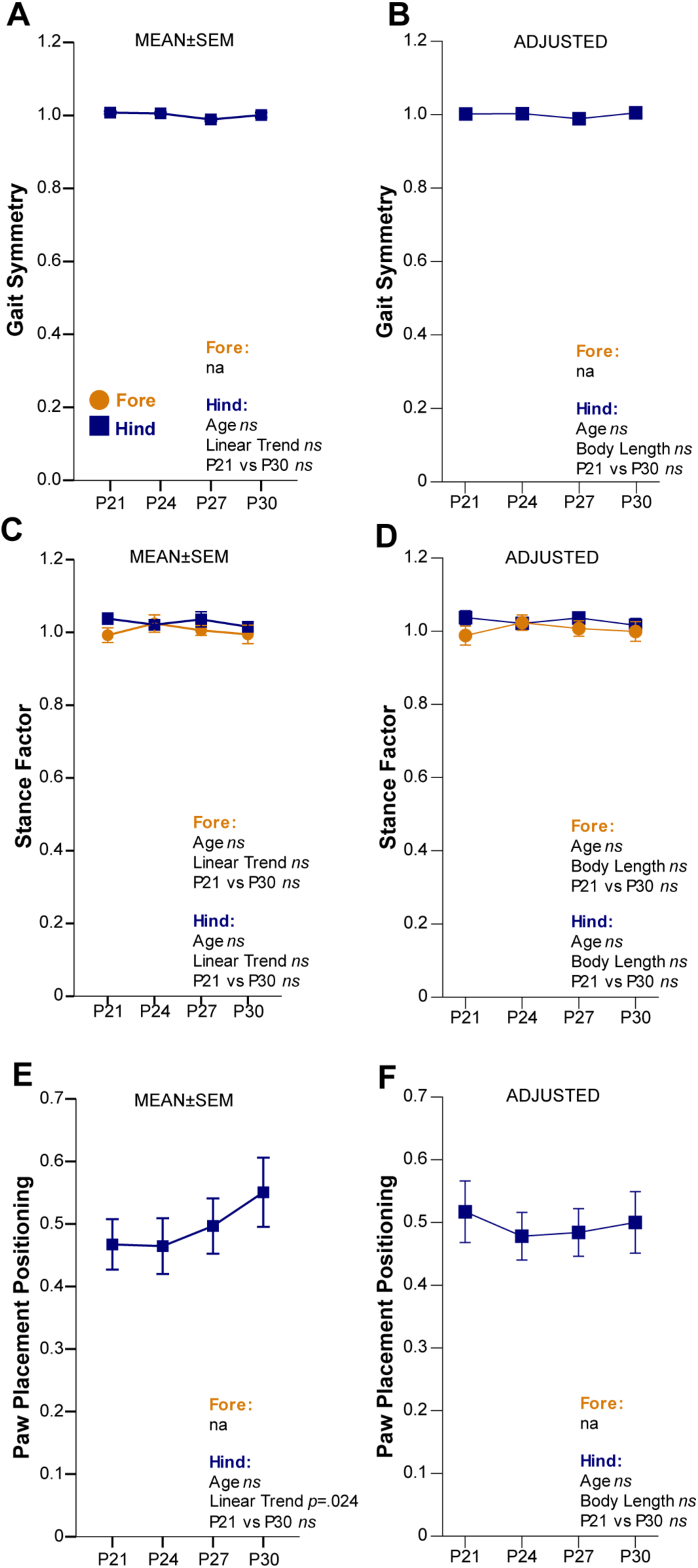
Gait metrics of balance and symmetry measured from P21 to P30 in C57 mice were not influenced by body length and did not change. (**A-F**) Gait symmetry, stance factor, and paw placement positioning means ± standard error of the mean (SEM) and covariate adjusted means (ADJUSTED) for body length (value 6.52078). For means ± SEM graphs, statistics derived from rmANOVAs include main effect of Age, Linear Trend Analysis, and *a priori* determined pairwise comparison of P21 to P30. For ADJUSTED graphs, statistics derived from LMM include main effect of Age, main effect of the covariate Body Length and *a priori* determined pairwise comparison of P21 to P30. A comprehensive list of statistical output can be found in Table 2.

### Spatiotemporal measurements of gait were influenced more by body length than age in FVB mice

In our FVB dataset, we again found many gait metrics that changed from P21 - P30 (Table 4), allowing for the identification of likely markers of gait maturation during this developmental window. The metrics influenced by age were primarily of the spatiotemporal subcomponent. No variability parameters and only one postural metric showed significant change with age. Body length significantly predicted 39% of the variables with a much less robust influence on direction of change for individual metrics compared to what we observed in the C57 dataset. Results from the rmANOVA suggested 22 of 44 metrics significantly changed from P21 to P30 (Table 3). However, after adjusting for body length in the LMM, this was reduced to 19 metrics, all of which were among the original 22. All metrics for FVB gait performance are discussed in detail below.

As in the C57 dataset, the important influence of body length was demonstrated by its role in stride frequency and stride length. Stride frequency and stride length appeared to decrease and increase, respectively, from P21 to P30 in the rmANOVA (Figure 7A,B). However, this was again due to the influence of body length. Body length was a significant predictor of both stride frequency and length for both fore- and hindlimbs (Figure 7C,D). While the LMM produced significant effects of age for both variables, the metrics fluctuated over time rather than following a consistent trajectory. In FVB mice the intra-step variability assessed by stride length CV was not influenced by body length nor age (Figure 7 E,F). This suggests, similarly to C57 mice, that stride lengths are already very consistent in FVB mice by P21.

**Figure 7.**
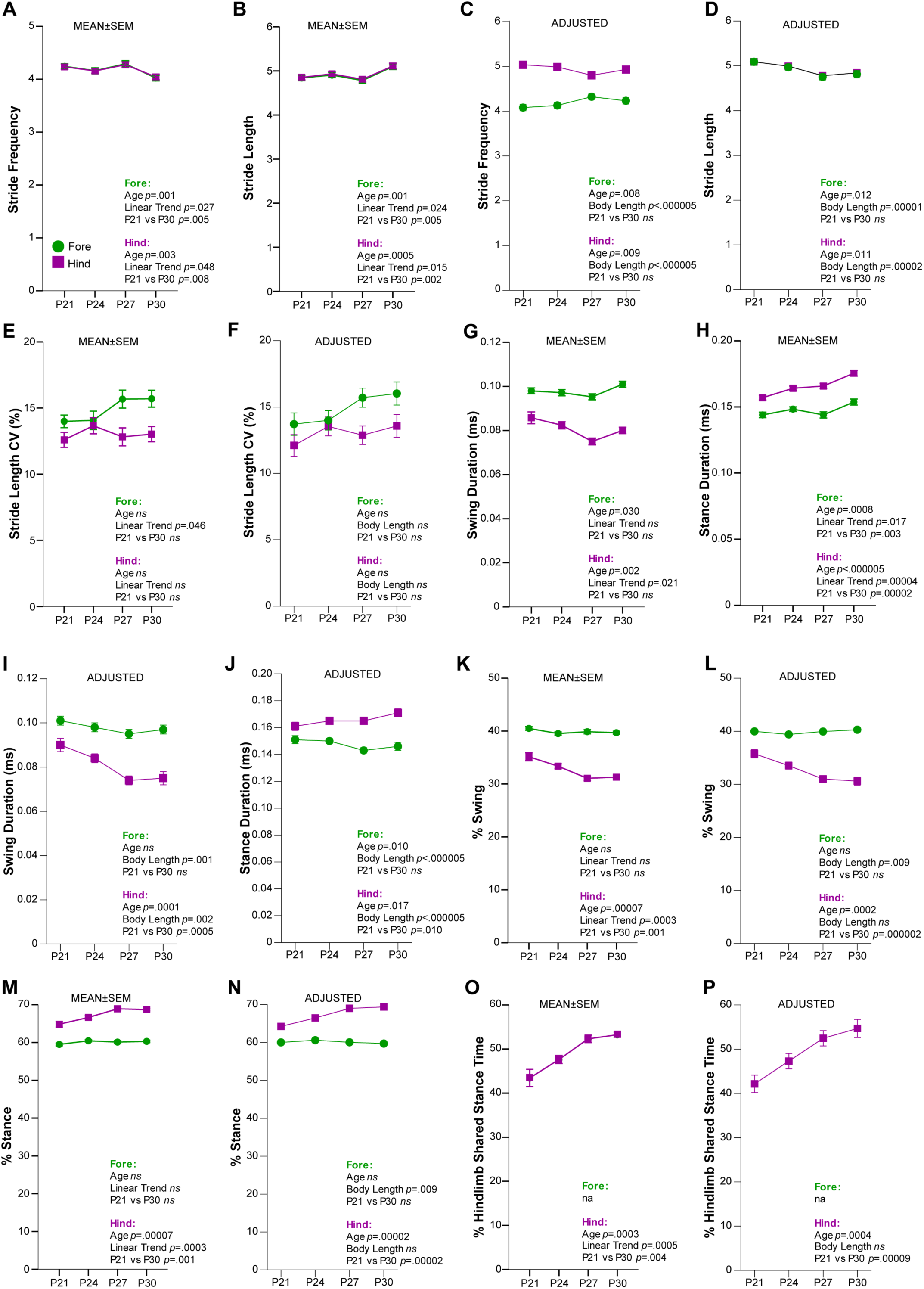
Main components of stride were significantly predicted by body length in FVB mice from P21 to P30. (**A-B**) Stride frequency and stride length means ± standard error of the mean (SEM). (**C-D**) Stride frequency and length covariate adjusted means (ADJUSTED) for body length (value 7.31226). (**E-F**) Stride length coefficient of variance (CV) means ± SEM and ADJUSTED. (**G-J**) Stance duration and swing duration means ± SEM and ADJUSTED. (**K-N**) Means ± SEM and ADJUSTED for %swing and %stance. (**O-P**) %Hindlimb Shared Stance Time means ± SEM and ADJUSTED. For means ± SEM graphs, statistics derived from rmANOVAs include main effect of Age, Linear Trend Analysis, and *a priori* determined pairwise comparison of P21 to P30. For ADJUSTED graphs, statistics derived from LMM include main effect of Age, main effect of the covariate Body Length and *a priori* determined pairwise comparison of P21 to P30. A comprehensive list of statistical output can be found in Table 3.

We next examined the two phases of stride: swing and stance. Results from the rmANOVA indicated no change in swing duration for either set of limbs and an increase in stance duration for both sets of limbs (Figure 7G,H). However, the LMM revealed body length as a significant predictor for swing duration and stance duration for both limbs (Figure 7I,J). Therefore, adjusting for body length changes over time revealed the hindlimbs change from P21 to P30 by decreasing time in swing and increasing time in stance. When we examined swing and stance as percentages of stride, body length was not a significant predictor of either %swing or %stance. Thus, the outcomes were the same for both the rmANOVA and LMM. For the forelimbs, the %swing was constant across age at 40% and the %stance was constant at 60%, reflective of the proportions observed in humans and P30 C57 mice (Figure 7K-N). For hindlimbs, we observed a significant decrease away from 40% for %swing and a significant increase away from 60% for %stance. These data suggest that FVB mice spend a greater proportion of their hindlimb strides in stance as they age over this developmental window.

The increases in stance duration and %stance were also reflected in the related spatiotemporal measurement, %hindlimb share stance time. This metric differed in the FVB dataset compared to what we observed in the C57 dataset. In FVB mice, body length did not significantly predict % of time in shared stance, and thus we saw a significant change from P21 to P30 in both the rmANOVA and LMM (Figure 7O,P). Also different from the C57 mice, FVB mice increased % time in shared stance whereas C57 remained unchanged across age for this metric. FVB mice have longer body lengths compared to C57 mice (Figure 1E). It is possible the change we see in %stance and %hindlimb shared stance reflect compensations for longer body size from P21 to P30 to remain at a speed of 20 cm/s - perhaps moving at the same speed with a longer body requires the FVB mice to spend increased time in stance.

The spatiotemporal metric paw overlap distance also shows an opposite pattern in the FVB data than we observed in the C57 data. Specifically, paw overlap distance significantly increased across age in FVB mice. While body length is a significant predictor of paw overlap distance, inclusion of body length as a covariate did not alter the direction of change across time. We saw the same significant increase from P21 to P30 in the rmANOVA and LMM analyses (Figure 8A,B). Like %hindlimb shared stance time, this increase in paw overlap distance may be a result of differences in body length (Figure 1E) and thus reflect compensatory mechanisms to remain at the same speed across regardless of size.

**Figure 8.**
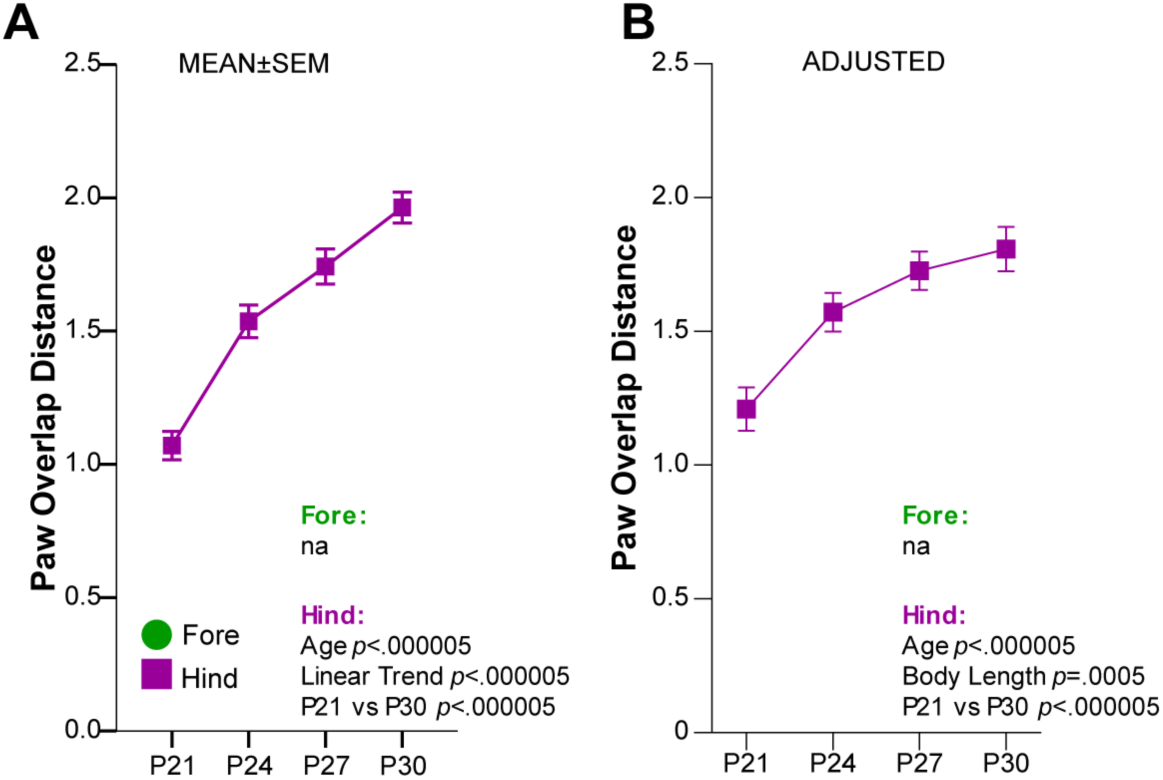
Paw overlap distance was significantly predicted by body length and increased from P21 to P30 in FVB mice. (**A-B**) Paw overlap distance means ± standard error of the mean (SEM) and covariate adjusted means (ADJUSTED) for body length (value 7.31226). For means ± SEM graphs, statistics derived from rmANOVAs include main effect of Age, Linear Trend Analysis, and *a priori* determined pairwise comparison of P21 to P30. For ADJUSTED graphs, statistics derived from LMM include main effect of Age, main effect of the covariate Body Length and *a priori* determined pairwise comparison of P21 to P30. A comprehensive list of statistical output can be found in Table 3.

### Gait maturation in FVB mice is reflected in components of the stance phase that differ from C57 mice

In C57 mice, we observed a balancing of forelimb and hindlimb contributions to both braking time and prolusion time as mice age (Figure 4A,B). In the FVB mice, the time the forelimbs spent in braking and propulsion remained unchanged from P21 to P30 (Figure 9A,B). The time the hindlimbs spent braking did increase with age, but the propulsion remained unaffected by age. As in the C57 dataset, body length was a significant predictor of both braking and propulsion times. Hindlimb propulsion appeared to significantly increase over time according to the rmANVOA results, however this was entirely due to the changes in body length (Figure 9B,C). The maximal rate of paw contact change, or time to load the paws onto the belt, for the braking phase increased for both forelimbs and hindlimbs, indicating that FVB mice load their paws onto the belt more quickly with age (Figure 9E,F). Peak paw area for the hindlimbs also increased with age (Figure 9G,H), however, paw length did as well, (Figure 9I), making it difficult to disentangle their respective contributions. Body length significantly predicted maximal rate of paw contact change in the hindlimbs not the forelimbs, therefore the results for the rmANVOA were similar to that for the LMM.

**Figure 9.**
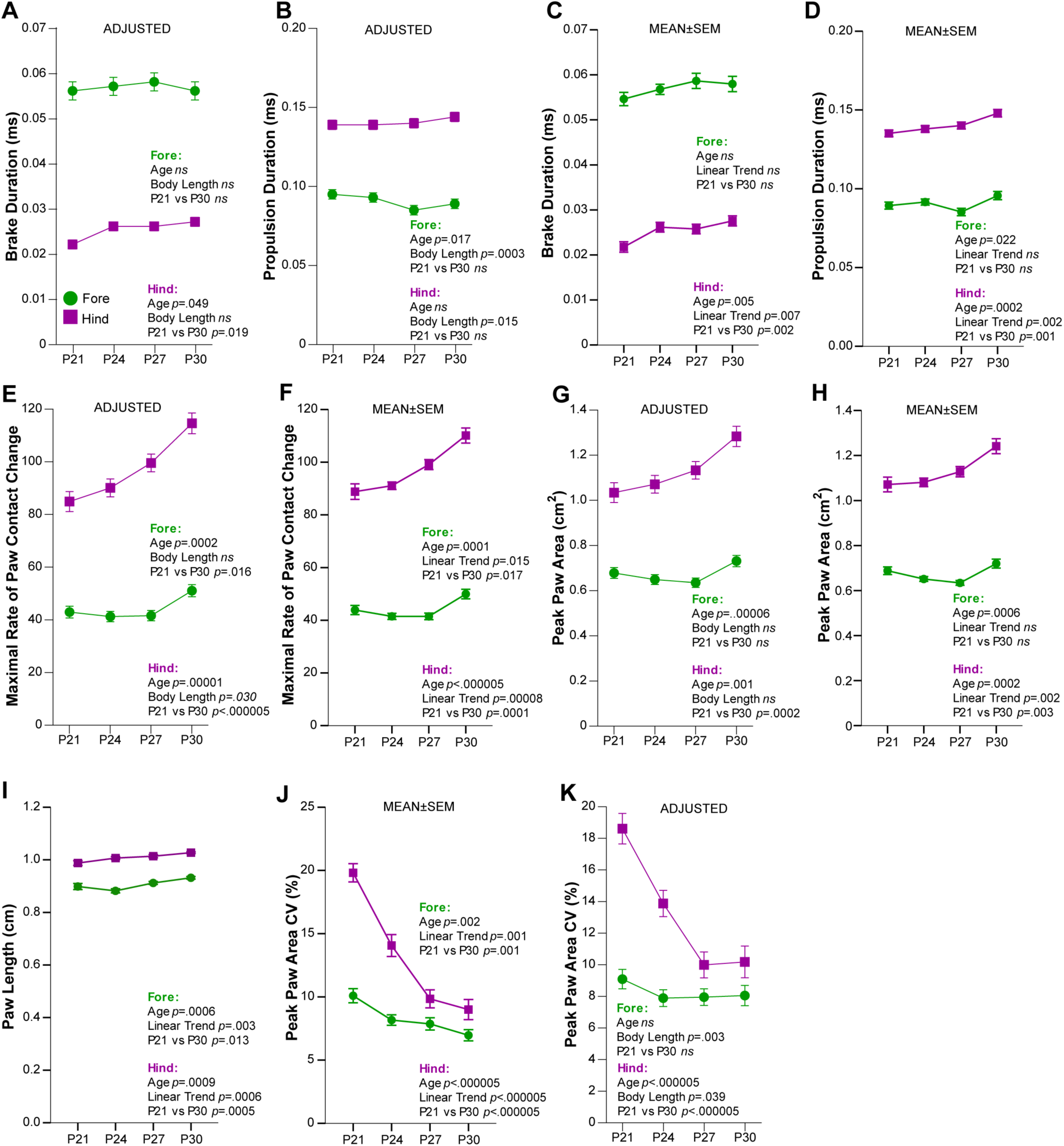
Hindlimb loading during the brake portion of the stance phase was dynamic from P21 to P30 in FVB mice. (**A-D**) Brake duration and propulsion duration covariate adjusted means (ADJUSTED) for body length (value 7.31226) and means ± standard error of the mean (SEM). (**E-F**) Maximal rate of paw contact change ADJUSTED and means ± SEM. (**G-H**) Peak paw area ADJUSTED and means ± SEM. (**I**) Paw length means ± SEM measured at full stance. (**J-K**) Peak paw area coefficient of variance (CV) means ± SEM and ADJUSTED. For means ± SEM graphs, statistics derived from rmANOVAs include main effect of Age, Linear Trend Analysis, and *a priori* determined pairwise comparison of P21 to P30. For ADJUSTED graphs, statistics derived from LMM include main effect of Age, main effect of the covariate Body Length and *a priori* determined pairwise comparison of P21 to P30. A comprehensive list of statistical output can be found in Table 3.

We observed a robust change in the variability of peak paw area (peak paw area CV) in these mice. The rmANOVA suggested peak paw area variability decreased for both forelimbs and hindlimbs decreased from P21 to P30 (Figure 9J). However, body length was a significant predictor of this metric, and the change in the forelimbs did not remain significant once body length was added as a covariate (Figure 9K). The variability in peak paw area in the hindlimbs at full stance robustly decreased. This may suggest the FVB mice were gaining greater control over paw placement during stance with age.

The remainder of the metrics analyzed were independent of both body length and age. This includes postural metrics, aspects of gait variability, and spatiotemporal measures of balance and gait symmetry, all of which we also found to be independent of body length and age in the C57 dataset, with the exception of stance width. Unlike in C57 mice, stance width remained unchanged across the examined developmental window in FVB mice (Figure 10A,B). The postural metrics absolute paw angle and step angle did not change with age nor were influenced by body length (Figure 10C-F). Almost all of the normalized variability metrics remained constant across this developmental window in FVB mice, excluding peak paw area CV described above. The CVs for stride length (as discussed above, Figure 7E,F), swing duration, stance width, paw angle, and step angle were all consistent from P21 to P30 and not significantly predicted by body length (Figure 10G-N). The measurements of balance and gait symmetry (gait symmetry, stance factor, and paw placement positioning, Figure 11A-F) were also independent of body length and remained unchanged across age. As suggested above, factors contributing to balance and symmetry likely develop at a much earlier age and are well established by the time a mouse is able to run on a treadmill system.

**Figure 10.**
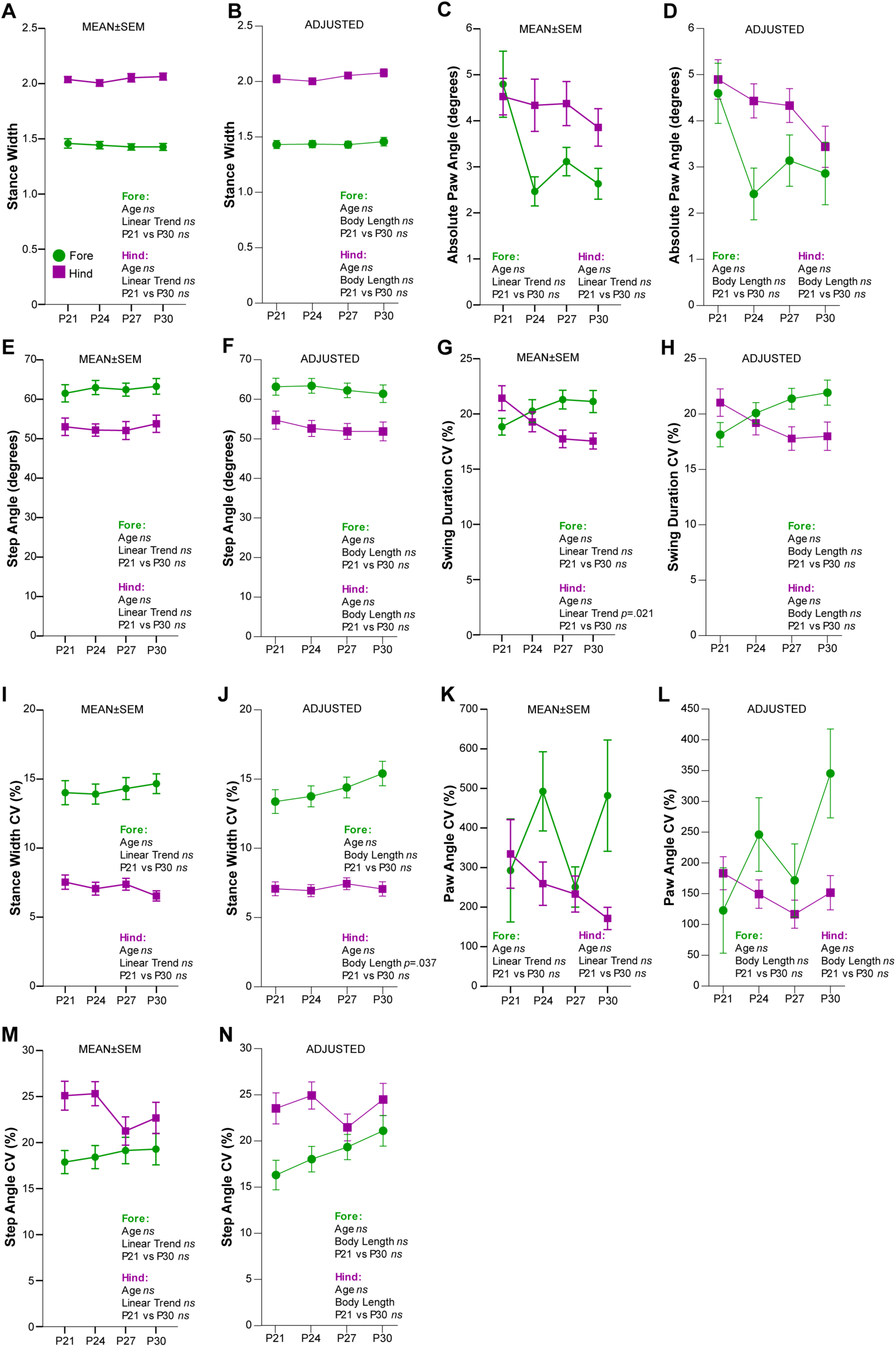
Stance width and most postural metrics and variability measures in FVB mice were independent of both body length and age. (**A-B**) Stance width means ± standard error of the mean (SEM) and covariate adjusted means (ADJUSTED) for body length (value 7.31226). (**C-F**) Absolute paw angle and step angle means ± SEM and ADJUSTED. (**G-J**) Swing duration coefficient of variance (CV) and stance width CV means ± SEM and ADJUSTED. **(K-N**) Paw angle CV and step angle CV means ± SEM and ADJUSTED. For means ± SEM graphs, statistics derived from rmANOVAs include main effect of Age, Linear Trend Analysis, and *a priori* determined pairwise comparison of P21 to P30. For ADJUSTED graphs, statistics derived from LMM include main effect of Age, main effect of the covariate Body Length and *a priori* determined pairwise comparison of P21 to P30. A comprehensive list of statistical output can be found in Table 3.

**Figure 11.**
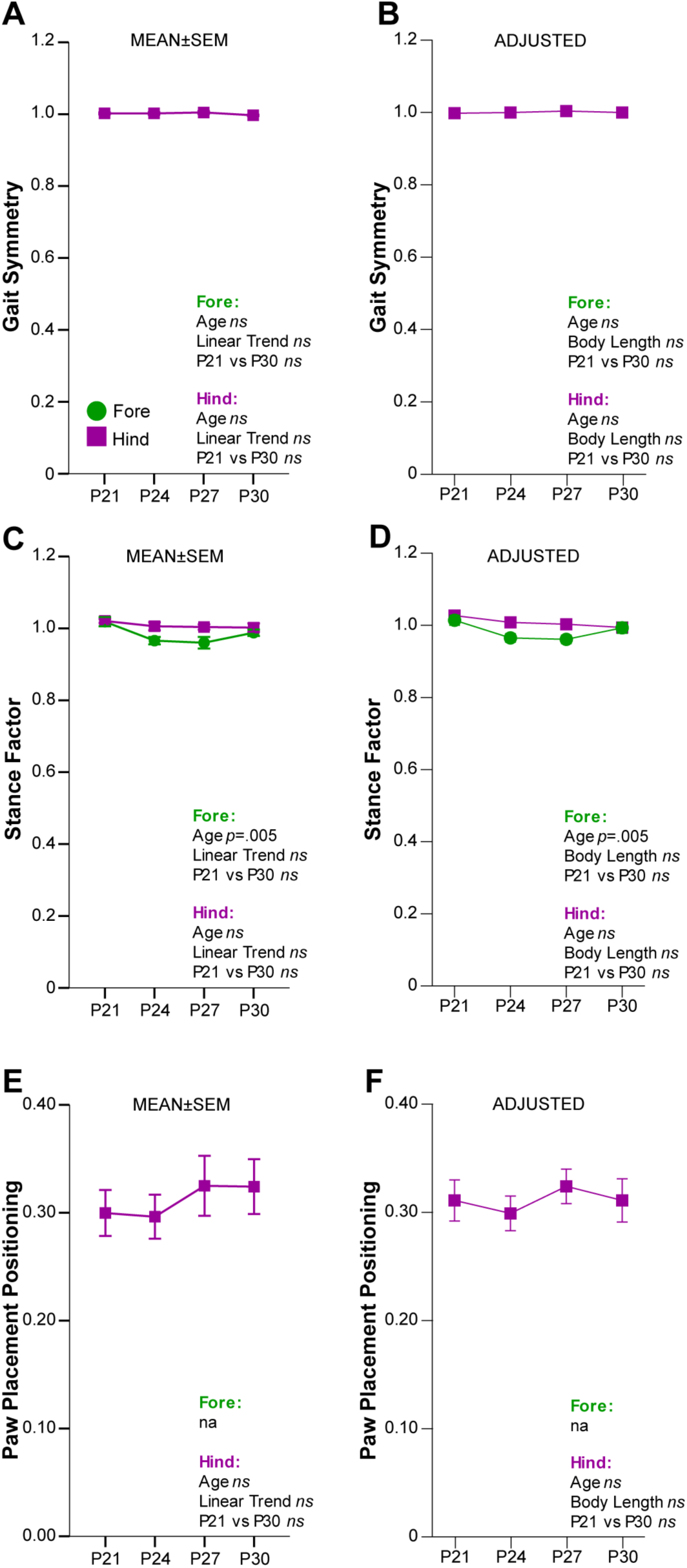
Gait metrics of balance and symmetry measured from P21 to P30 in FVB mice were independent of both body length and age. (**A-F**) Gait symmetry, stance factor, and paw placement positioning means ± standard error of the mean (SEM) and covariate adjusted means (ADJUSTED) for body length (value 7.31226). For means ± SEM graphs, statistics derived from rmANOVAs include main effect of Age, Linear Trend Analysis, and *a priori* determined pairwise comparison of P21 to P30. For ADJUSTED graphs, statistics derived from LMM include main effect of Age, main effect of the covariate Body Length and *a priori* determined pairwise comparison of P21 to P30. A comprehensive list of statistical output can be found in Table 3.

## Discussion

Gait disruptions can represent a pathological state across many neurological diseases and disorders. These abnormalities can reflect a breakdown of motor control circuits as in neurodegenerative diseases like Parkinson’s Disease and Huntington’s Disease (Amende et al., 2005; Hausdorff et al., 1998; Karachi et al., 2010; Laforet et al., 2001; Rao et al., 2008), or a faulty maturation of motor control circuits as in neurodevelopmental disorders (Jeste, 2011; Mosconi et al., 2015). Here, we have quantitatively characterized typical development of multiple gait parameters in C57 and FVB mice in a controlled setting, accounting for major confounding factors in gait analysis.

Examination of gait in these two oft-used mouse strains revealed a set of gait metrics that change over the P21-P30 developmental period, and thus likely reflect aspects of gait maturation. In C57 mice, we observed a decrease in stance duration overall, and a balancing of the phases of stance between limbs. At P21 there was a robust difference in the time spent in both braking and propulsion by each limb, but by P30 each limb spent equal time in braking and propulsion. We also observed a decrease in the rate at which the hindlimbs were loaded into the stance phase, which is likely related to the observed increased duration of hindlimb braking. Finally, the peak area of the paw at full stance decreased with age, which may reflect a change from a flat-footed stance to heel-to-toe stance. Peak paw area may be a valuable parameter to investigate the role of paw and toes in gait, such as toe walking, which is common in NDDs.

In FVB mice, aspects of gait maturity were reflected in stance-related metrics as well, albeit with different trajectories than those seen in C57 mice. FVB mice increased braking time with their hindlimbs, similar to C57s, yet they also increased their rate of deceleration or loading of paws into stance. They exhibited an increase in peak paw area over time, opposite to that observed in C57 mice. As noted above, it is difficult to separate this change from increased paw size and thus it remains uncertain how this finding relates to gait maturation. What is perhaps more clearly interpretable is the decrease in variability exhibited by FVB mice in their peak paw area over time - this may reflect more precise placement of the paw during the stance phase as the mice age. Across both strains, the most sensitive measures of gait maturation during the P21-P30 developmental window are these distinct elements of stance.

Several important methodological considerations distinguish this study from prior work in the literature. Most studies of gait in mouse models of disease, including NDDs, are conducted in adult animals (Amende et al., 2005; Gadalla et al., 2014; Galante et al., 2009; Kloth et al., 2015; Schneider et al., 2012). While this is a useful endeavor to understand gait abnormalities in the mature animal, these studies provide no information about the development of such abnormalities of how they might vary over time. The few studies that have examined mouse gait at earlier time points provide valuable information at specific ages (Wozniak et al., 2019), but represent only a snapshot of gait performance in time. In contrast, here we have characterized gait across multiple timepoints in a longitudinal manner and at a consistent speed, enabling the sensitivity of a repeated measures design and allowing accurate comparisons of gait across age. This baseline characterization of healthy gait development will inform future studies of NDD models, providing insight into how and when gait irregularities arise across development and whether these represent delays in normal development, or completely distinct trajectories.

We also recognized that the rapid growth in body size across the age range examined could confound our gait data, and indeed we found that more than a third of all metrics were significantly influenced by body length. Decreased stride frequency, or cadence, and increased stride length have been suggested as markers of gait maturity in humans, however they are mainly driven by limb lengthening and thus likely do not reflect underlying motor circuit maturation - a more relevant phenotype for brain disorders like NDD. In both strains examined, we initially observed a comparable pattern of decreased stride frequency and increased stride length. However, body length significantly impacted both of these variables, and controlling for this influence revealed that neither stride frequency nor length changed with age from P21 to P30. Thus, we presented the data both before and after controlling for the impact of body size on each gait parameter to highlight the possibility of erroneous interpretations when body size is not considered, and to help define those features that could reflect true differences in CNS circuits rather than simply changes in limb length.

Understanding the development of mouse gait is most useful if it can inform the consequences of homologous genetic lesions seen in humans. The gait analysis system used here provides a comprehensive set of gait metrics, most of which have not been examined across development in humans and thus we cannot comment on their translational impact at this time. However, we did find key parallels in our results to that which has been observed in human gait development. Specifically, the stride of our mice at P30 was composed of 40% swing phase and 60% stance phase, mirroring the mature human stride composition (Hillman et al., 2009; Lythgo et al., 2011; Sutherland et al., 1980). There was some drift in the hindlimbs of FVB mice; however, this may be a compensatory mechanism to allow the animal maintain a consistent speed. Nonetheless, these findings suggest the translational potential of this approach to interrogate the impact of mutations on gait circuitry in mouse models of human disease.

Our study had several limitations, some of which were trade-offs intended to maximize consistency between ages tested. For example, the mice were limited to a forced speed across all four timepoints. As body size increases with age, the intrinsic speed or qualitative gait type at a given speed (e.g. trot versus run) may change as well. This could influence the change, or lack thereof, of some of the variables we measured. However, as speed is the greatest modulator of gait, appropriate comparisons required us to enforce a constant speed for all mice across all ages. Regardless of the variation in gait that might be revealed at different speeds, this study provides a benchmark of gait at 20 cm/s across the developmental window from P21-P30, defining an assay that will be valuable as we begin to study how genetic and environmental disruptions of neurodevelopment impact gait development.

While different strains of mice often show differential performance in in many behavioral assays (Eisener-Dorman et al., 2011; Keum et al., 2016; Liu et al., 2011; Martin et al., 2014; Moy et al., 2004, 2008), our study was not designed to directly compare the two strains tested so our ability interpret any differences is limited. The differences observed may arise from size discrepancies, as FVB mice are consistently larger, or experimental parameters, such as different sample sizes. It is also possible that motor development in the FVB mouse occurs earlier than in the C57 mouse, as suggested by the fact that FVB mice were able to run successfully on the DigiGait at a younger age (P17) than C57 mice (P21, pilot data not shown). Future direct comparison studies are needed to validate strain differences suggested by the present study. In any case, our results provide a reference dataset of gait maturation for two commonly used mouse strains, often employed as the genetic backgrounds for models of disease.

Ultimately, the results of this study are a normative standard against which murine models of NDDs may be compared. Mouse models of NDDs are inherently limited due to the primary cognitive impairments in these disorders often being of processes specific to humans, and only paralleled in mice. However, gait abnormalities and changes in gait development are some of the few features of NDDs that may track from murine models to human disease phenotypes, as neural control of gait has many shared neural mechanisms between mouse and human (Dominici et al., 2011; Takakusaki et al., 2008). Thus, this study provides the foundation for future phenotyping of gait in mouse models that will serve as a vital window into understanding the disruption of motor circuits in human disease.

## Author Contributions Statement

Data collection was conducted by SA, KW and CW. Data analysis was conducted by SM. Manuscript writing was done by SA, KM, CW and SM. SA, JD and SM edited the manuscript and designed the study.

## Acknowledgements

The authors thank Michael Vasek, PhD for valuable discussion during the preparation of this manuscript and Rena Silverman for aiding us with paw measurements. In addition, the authors thank the Co-Directors of the Animal Behavior Core at Washington University School of Medicine, David Wozniak, PhD and Carla Yuede PhD, as well as Azad Bonni, MD, PhD, for access to the DigiGait equipment.

## Supplementary Material

### 1. Supplementary Figures

**Supplementary Figure 1.**
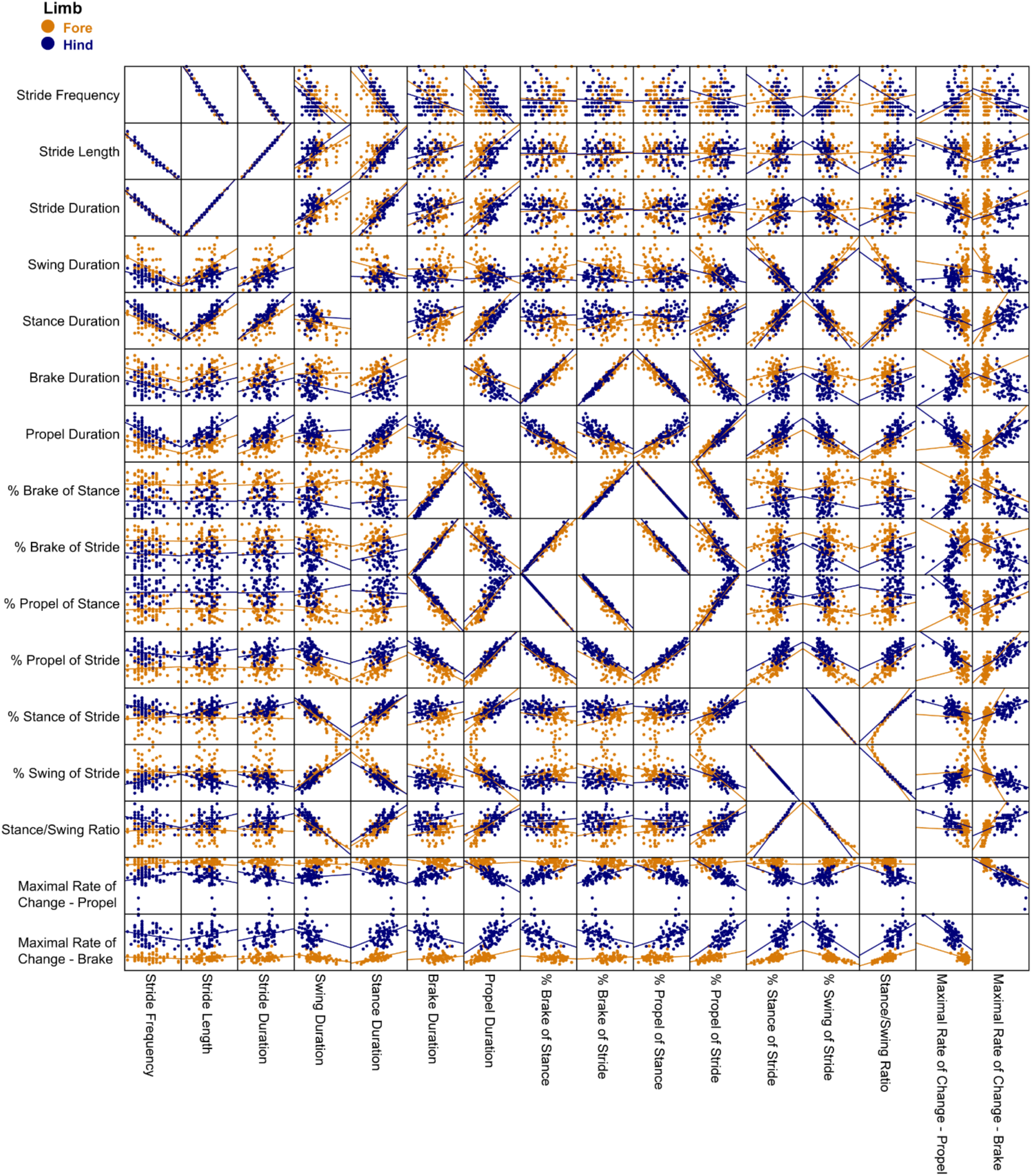
Scatter matrices of spatiotemporal metrics measured in both fore and hindlimbs of C57 mice at P21, P24, P27 and P30. Pairs of variables that were considered close to or at perfect alignment are outlined in red boxes. The variable that was excluded from further analysis is also outlined.

**Supplementary Figure 2.**
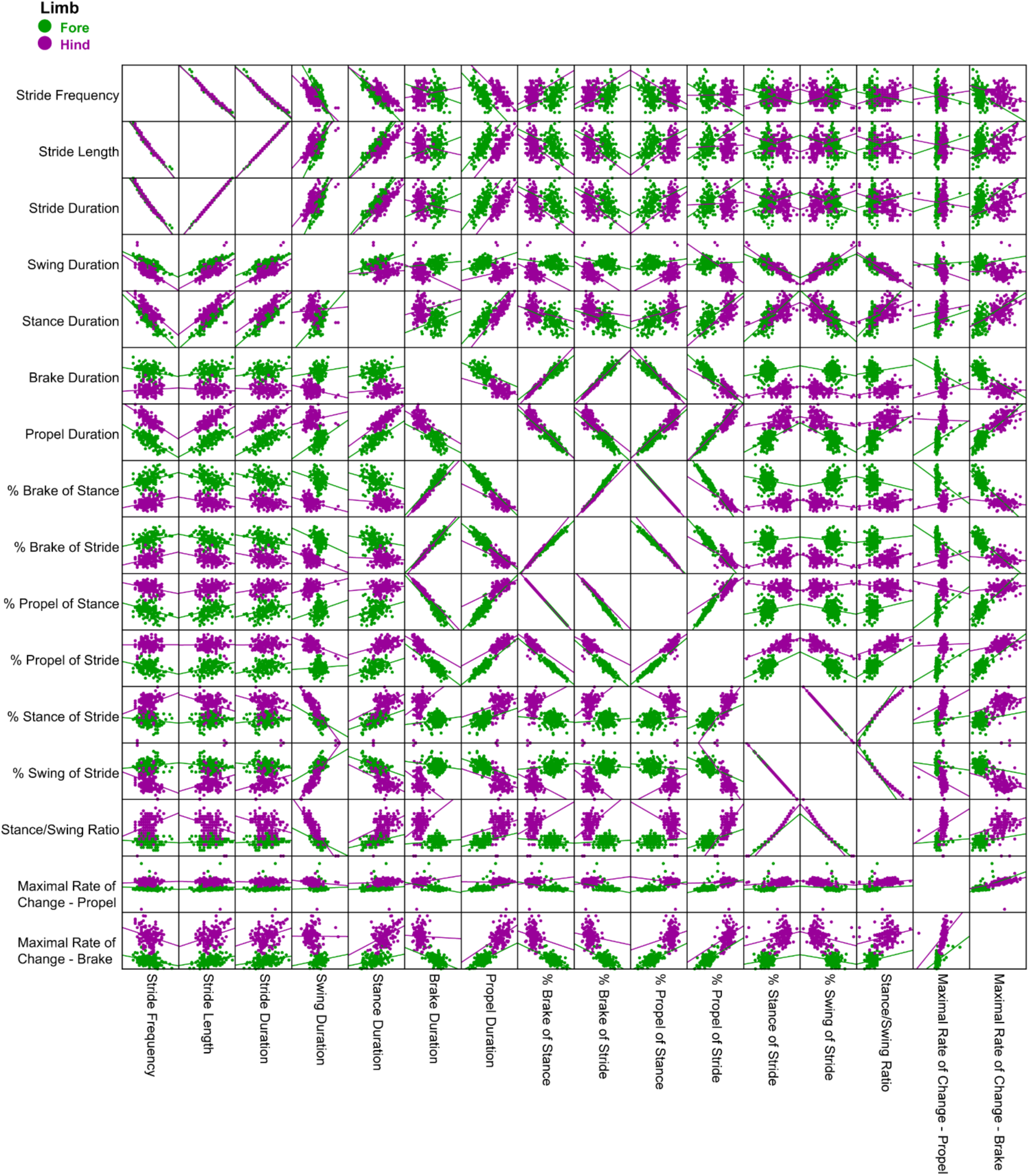
Scatter matrices of spatiotemporal metrics measured in both fore and hindlimbs of FVB mice at P21, P24, P27 and P30. Pairs of variables that were considered close to or at perfect alignment are outlined in red boxes. The variable that was excluded from further analysis is also outlined.

**Supplementary Figure 3.**
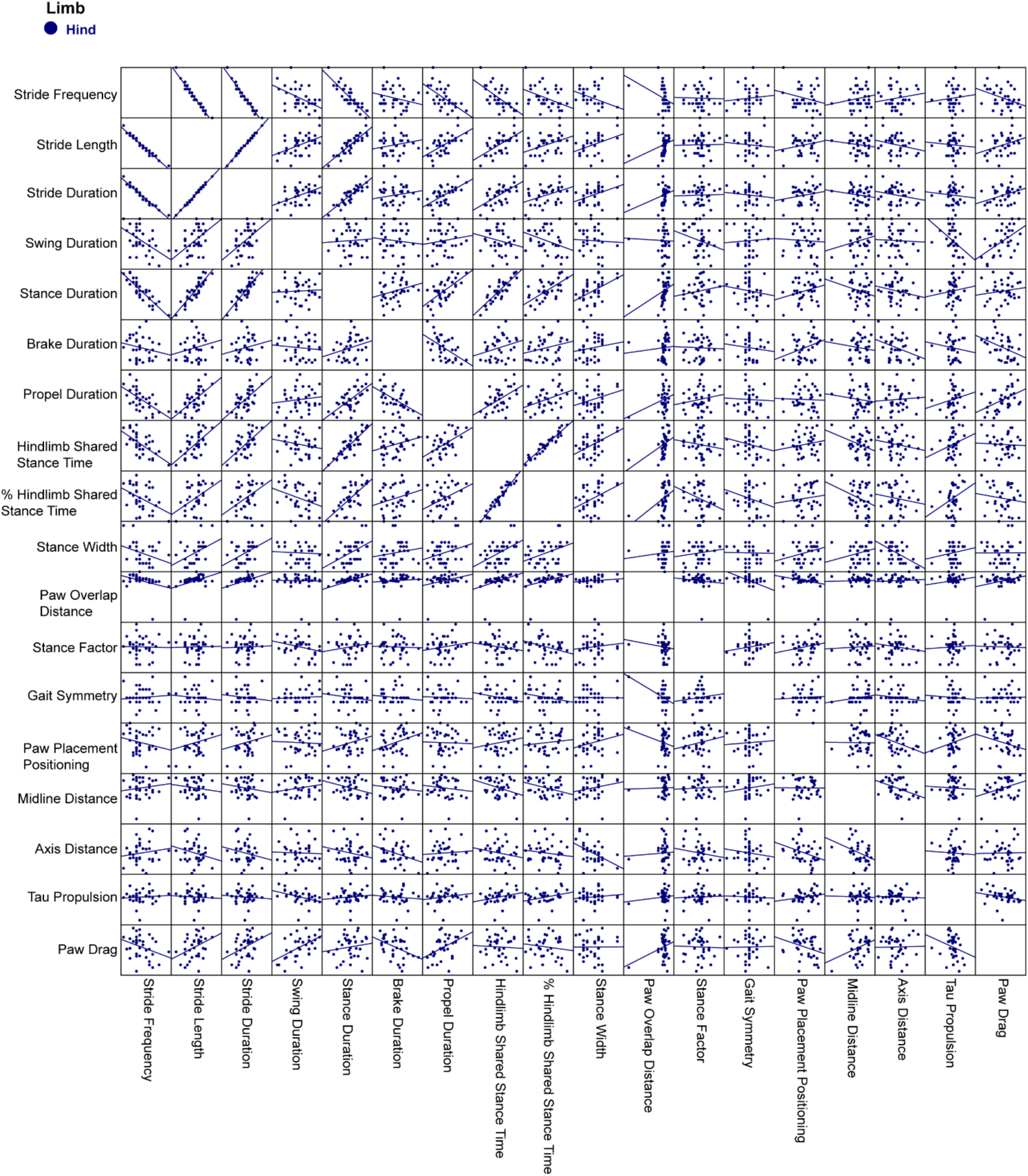
Scatter matrices of spatiotemporal metrics measured in only the hindlimbs of C57 mice at P21, P24, P27 and P30. Pairs of variables that were considered close to or at perfect alignment are outlined in red boxes. The variable that was excluded from further analysis is also outlined.

**Supplementary Figure 4.**
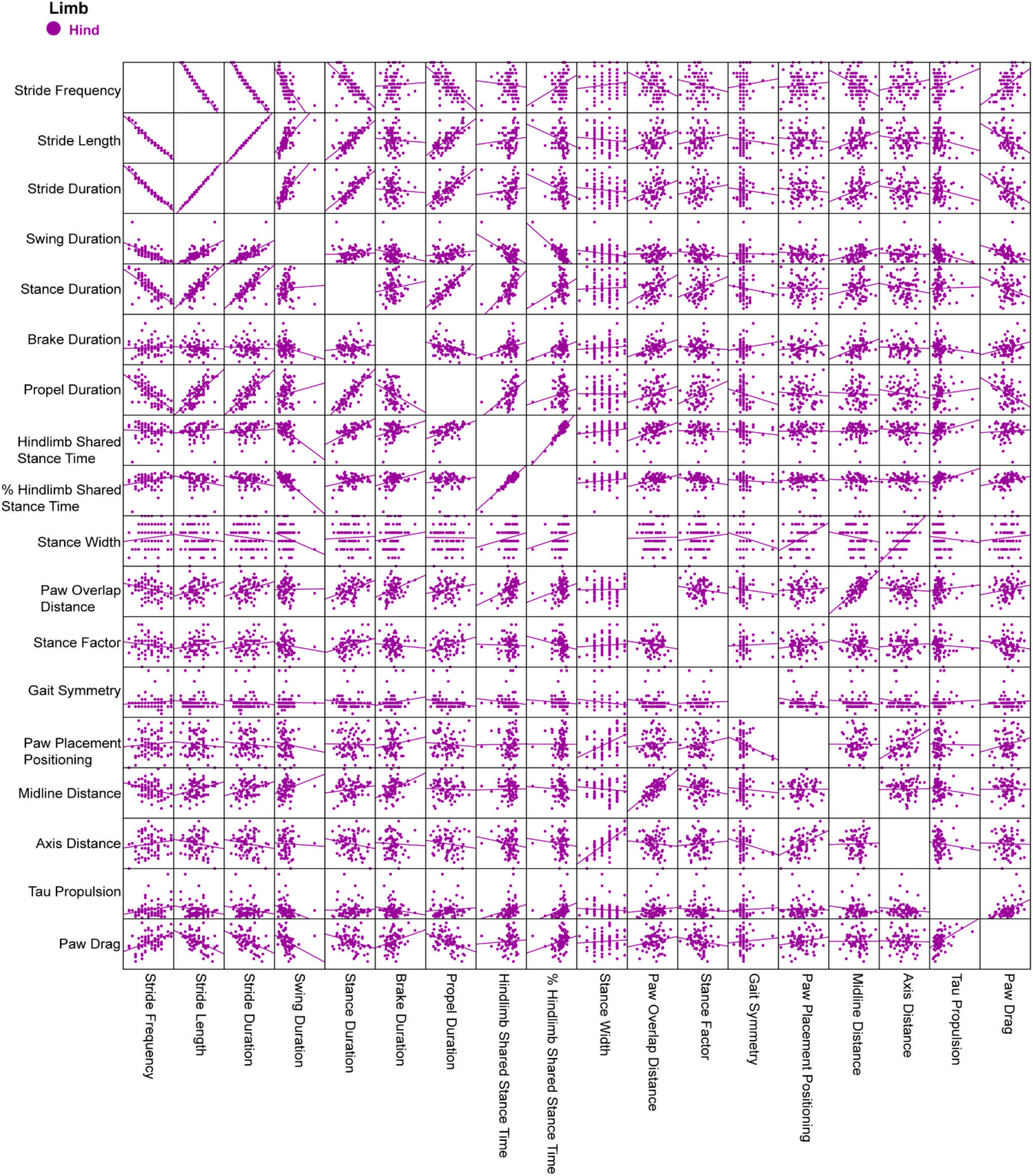
Scatter matrices of spatiotemporal metrics measured in only the hindlimbs of FVB mice at P21, P24, P27 and P30. Pairs of variables that were considered close to or at perfect alignment are outlined in red boxes. The variable that was excluded from further analysis is also outlined.

**Supplementary Figure 5.**
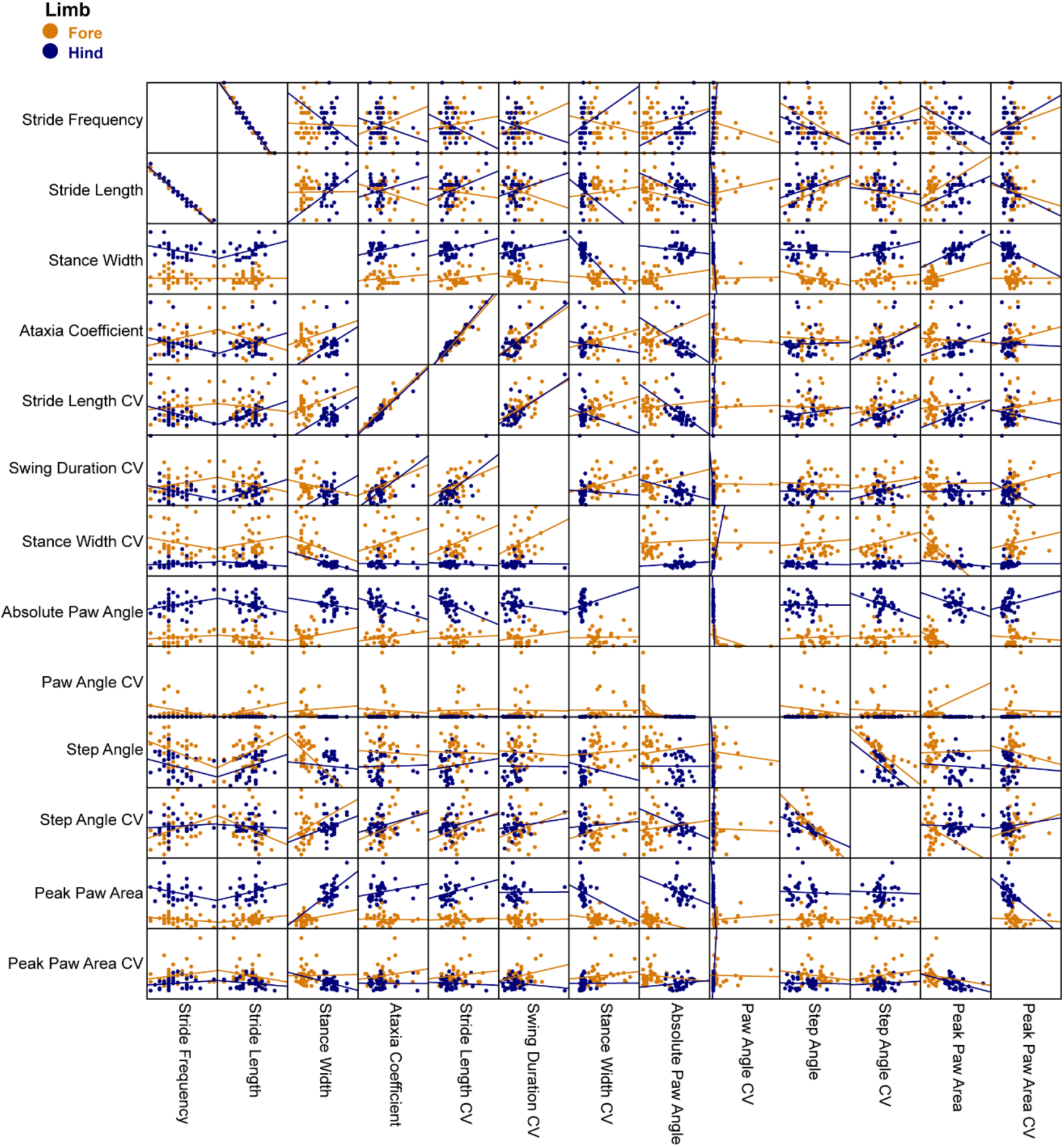
Scatter matrices of postural and variability metrics measured in both fore and hindlimbs of C57 mice at P21, P24, P27 and P30. Pairs of variables that were considered close to or at perfect alignment are outlined in red boxes. The variable that was excluded from further analysis is also outlined.

**Supplementary Figure 6.**
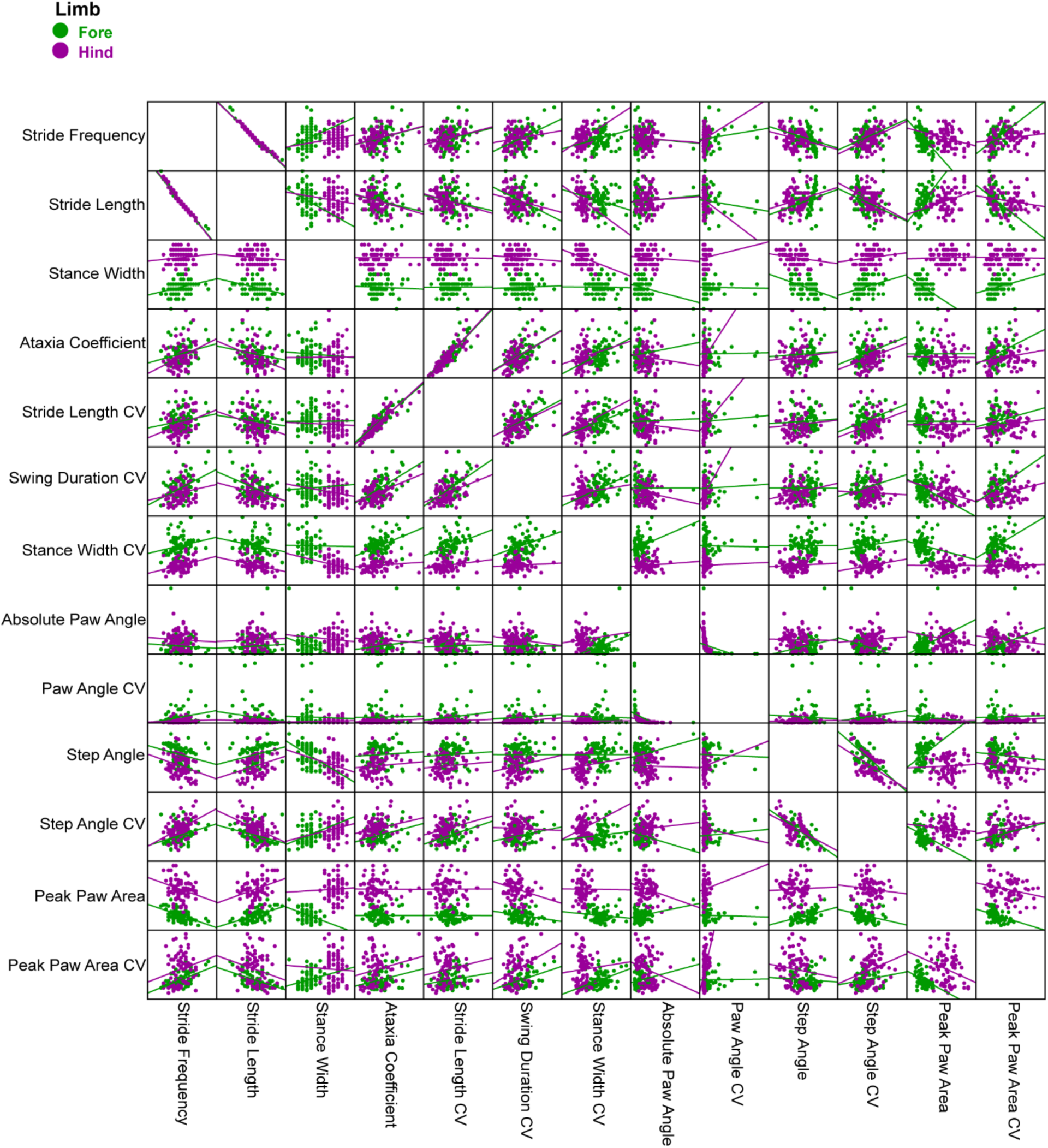
Scatter matrices of Scatter matrices of postural and variability metrics measured in both fore and hindlimbs of FVB mice at P21, P24, P27 and P30. Pairs of variables that were considered close to or at perfect alignment are outlined in red boxes. The variable that was excluded from further analysis is also outlined.

